# A splenic reservoir of pro-angiogenic neutrophils supports blood flow recovery after peripheral tissue ischemia

**DOI:** 10.64898/2026.10.21.760850

**Authors:** Catarina Amoedo-Leite, Kristel Parv, Cédric Seignez, Anna Seidler, Hanna Ermesjö, Olha Pavliuk, Carlota F Prendes, Kevin Mani, Robin Lindsay, Gustaf Christoffersson, Anders Wanhainen, Mia Phillipson

## Abstract

**Background:** Neutrophils accumulate at sites of ischemic injury, where a distinct subset characterized by CD49d^+^CXCR4^hi^VEGFR1^hi^ expression promotes angiogenesis and contributes to blood flow recovery. This study investigated the function, origin, and recruitment mechanisms of these pro-angiogenic neutrophils (PANs) in a murine model of hindlimb ischemia and explored whether a corresponding population responds to acute ischemia in humans.

**Methods:** Using spectral flow cytometry and immunohistochemistry, PANs and classical neutrophils were characterized in ischemic gastrocnemius muscle following hindlimb ischemia in mice. Blood flow recovery was assessed by laser speckle flowmetry. To determine the origin and mobilization mechanisms of PANs, splenectomy, chemical sympathectomy with 6-hydroxydopamine (6-OHDA), and surgical splenic denervation were performed. Translational studies examined blood samples collected during elective abdominal aortic aneurysm repair to assess PAN mobilization following acute lower limb ischemia induced by aortic cross-clamping.

**Results:** PANs rapidly accumulated in ischemic muscle through a CD49d-dependent mechanism, and blockade of CD49d impaired blood flow recovery. During homeostasis, PANs were maintained within a splenic reservoir and were rapidly released into circulation following ischemia. Splenectomy and disruption of splenic sympathetic signaling significantly reduced PAN recruitment to ischemic tissue and impaired functional recovery. Mobilization was driven by ischemia-induced sympathetic activation, which decreased CXCL12 levels in the splenic red pulp, disrupting CXCR4-mediated PAN retention. In humans, a phenotypically similar PAN population was identified in peripheral blood, which increased rapidly following lower limb ischemia during aortic cross-clamping. The speed of this response is consistent with release from an extramedullary reservoir, although its anatomical source remains unknown.

**Conclusion:** These findings identify the spleen as a reservoir of pro-angiogenic neutrophils in mice and reveal a neuroimmune mechanism regulating their mobilization and recruitment after ischemic injury. The rapid appearance of PAN in human circulation after acute ischemia suggests that comparable extramedullary neutrophil reservoirs may also exist in humans.

## Introduction

Tissue ischemia results from insufficient blood flow and oxygen delivery to meet metabolic demands, causing cellular injury that can progress to irreversible tissue damage and organ dysfunction if perfusion is not restored.^1,2^ Minor ischemic injuries are resolved through coordinated inflammatory and angiogenic responses that promote vascular repair and blood flow restoration through the release of pro-angiogenic mediators such as vascular endothelial growth factor (VEGF), endothelial cell activation, and immune cell recruitment.^2–4^ However, when ischemic injury is extensive or prolonged, these endogenous repair mechanisms may be insufficient to prevent irreversible tissue damage. Ischemic conditions, including myocardial infarction, stroke, and critical limb ischemia, contribute substantially to global morbidity and mortality.^5,6^ Current treatment strategies aim to restore tissue perfusion through pharmacological interventions, thrombolysis, angioplasty, stenting and bypass surgery. Nevertheless, many patients experience incomplete recovery despite restoration of large vessel blood flow, partly due to persistent inflammation, microvascular dysfunction, and inadequate vascular repair.^7,8^ Consequently, therapeutic angiogenesis has emerged as a promising strategy to improve tissue recovery and functional outcomes following ischemic injury, although its clinical efficacy remains to be demonstrated.^3,9^ Understanding the endogenous mechanisms that promote vascular repair may therefore provide new opportunities for therapeutic intervention.

Neutrophils are the most abundant circulating leukocytes in humans and among the first leukocytes to accumulate at the injured site following a tightly coordinated recruitment cascade including endothelial cell activation and chemokine signaling.^3,10–15^ They are traditionally recognized for their important roles in host defence and acute inflammation, but are increasingly acknowledged also for their contributions to tissue repair.^4,15,16^ At sites of injury, neutrophils have been shown to contribute to vascular remodelling and resolution of inflammation through the production of growth factors, clearance of cellular debris, and modulation of macrophage polarization.^3,17^ In fact, a distinct neutrophil population named pro-angiogenic neutrophils (PANs) has been identified that directly support angiogenesis and vascular remodelling.^3,18–20^ PANs are characterized by expression of CD49d, VEGFR1, and high levels of CXCR4, and represent only approximately 1–3% of circulating neutrophils in mice and humans during homeostasis.^18,20^ They are selectively recruited to ischemic tissues in a VEGF-A-dependent manner, where they support neovascularization and vascular remodelling through the release of matrix metalloproteinase 9.^20^ We previously demonstrated that recruitment of PANs is required for efficient angiogenesis and revascularization following syngeneic transplantation of pancreatic islets into skeletal muscle, as inhibition of PAN trafficking impaired neovessel formation and graft vascularization.^18,20^ Pro-angiogenic neutrophils have also been reported in experimental tumour models, suggesting that the capacity of neutrophils to regulate vascular growth extends across multiple pathological settings.^21,22^

Given their low frequency in the circulation, important unanswered questions are how sufficient numbers of PANs can rapidly be mobilized to ischemic tissues, and from where these cells originate. While neutrophils are generated in the bone marrow through granulopoiesis, they can also be maintained in extramedullary reservoirs, including the spleen and lungs, which may provide readily mobilizable populations during inflammation.^12,23–26^ Whether specialized neutrophil subsets such as PANs are preferentially maintained in such reservoirs, and how their release is regulated in response to ischemic injury, remains poorly understood.

Here, we investigated the functional role, origin, and mobilization of PANs in response to large-scale ischemia using a mouse model of hindlimb ischemia (HLI). We hypothesized that PANs contribute to vascular recovery following ischemic injury and sought to identify their source and the mechanisms governing their mobilization and recruitment. To assess the relevance of these findings in humans, we further investigated the response of a corresponding PAN population to acute lower-limb ischemia induced by temporary aortic cross-clamping during open abdominal aortic aneurysm (AAA) repair.

## Materials and methods

### Mice

Male C57Bl/6J mice (27.0 ± 0.5g, from Taconic Bomholt, Ry, Denmark) were used in the experiments, and were housed in the animal facility of Uppsala University Biomedical center until the start of the experiment. All mice had *ad libitum* access to food and water, and all experiments were approved by the Uppsala Region Laboratory Animal Ethics Board.

### Hindlimb ischemia model and treatments

Induction of hindlimb ischemia (HLI) was carried out as previously described.^27^ Briefly, mice were anesthetized (Isoflurane, 2-3%, Forene, Abbott Laboratories) and the fur was shaved from the left hindlimb. Animals received preoperative buprenorphine 0.5 mg/kg subcutaneously. Following opening of the skin, the femoral artery was separated from the nerve and femoral vein, followed by ligation and cutting of the artery between proximal caudal femoral artery and popliteal artery. After surgery, mice were allowed to wake up under a heat lamp and received buprenorphine 0.5 mg/kg subcutaneously 24h after surgery. Laser Speckle flowmetry was carried out the next day to confirm ischemia. Inhibition of CD49d was achieved by injection of a selective inhibitor of CD49d (#BIO5192; Tocris) (1 mg/kg of body weight in 200 µl of saline solution) both intravenously (i.v.) and intraperitoneally (i.p.) starting one day prior to onset of HLI and then twice daily until day 7. Systemic sympathectomy was achieved by intraperitoneal injection of 6-OHDA every 48 hours, starting 7 days before HLI induction. Control mice received an intraperitoneal injection of ascorbic acid, the vehicle of the 6-OHDA injection, every 48 hours, starting 7 days before HLI induction. All mice were euthanized by cervical dislocation at end of experiment.

### Laser Speckle assessment of functional tissue perfusion

Blood flow in footpads was measured using an in-house developed model where anesthetized mice were placed in prone position with paws resting on tubing with circulating water using Perimed Laser Speckle Contrast Analysis method (785 nm laser, 20 µm resolution) ^28^. After baseline registration (2 min, 22°C), the circulating water temperature was increased, resulting in a 10°C temperature increase on the skin of the footpad. Once blood flow measurement had stabilized, it was recorded for 2 min at 10 images/s and analyzed in PIMSoft (Perimed).

### Splenectomy

Splenectomy was performed on anesthetized mice (Isoflurane, 2-3%, Forene, Abbott Laboratories) as described before.^29^ Animals received preoperative buprenorphine 0.5 mg/kg subcutaneously. Briefly, laparotomy incision was made, and the splenic vein and artery were identified, ligated with a 7-0 silk suture and sectioned, and the spleen was removed. The laparotomy incision was closed using 6-0 silk suture. After surgery, the mice woke up under a heat lamp and received buprenorphine 0.5 mg/kg subcutaneously 24h after surgery.

### Splenic denervation

Splenic denervation was performed and confirmed by Tyrosine hydroxylase (TH) staining, as described in Zhang et al., 2020.^30^ After the mice were anesthetized, the peritoneal cavity was accessed through a midline abdominal incision. The spleen was isolated from the peritoneal cavity, and the three main supplying vascular trees were exposed. Absolute ethanol was repeatedly applied to vascular trees to deplete the splenic nerve fibres, and vasculature was evaluated for signs of vasoconstriction, taking breaks between ethanol applications to avoid damage to the vascular trees. For sham-operated mice, surgery was performed identically; however, a saline solution was used instead of absolute ethanol. Spleen weight and morphology was compared between groups to access for signs of autosplenectomy or splenic atrophy.

### Single cell isolation and flow cytometry

Mice were anesthetized (Isoflurane, as described above), and cardiac puncture was carried out to collect blood. After cervical dislocation, bone marrow, lung, spleen and gastrocnemius muscle were collected. The tissues were dissociated mechanically and enzymatically as follows: The muscle and lung were incubated in collagenase II (1 mg/ml) (#17101015; Gibco) in RPMI1640 in 37 degrees for 30 minutes. After washing with ice-cold DPBS, the muscle was incubated for additional 30 minutes with collagenase IV (0.5 mg/ml) (#17104019; Gibco) and dispase (2.5 mg/ml) (#17105041; Gibco) in RPMI1640, with intermittent pipetting. Thereafter, tissue suspension was passed through a 23G 0,6 mm needle 8-10 times, and filtered through a 70 µm cell strainer. The spleen was dissociated by passing through a 70 µm cell strainer. Red blood cell lysis was carried out using ACK lysing buffer (#A1049201; Gibco). Cell count was carried out using Invitrogen Countess II Automated Cell counter from Thermo Fisher Scientific.

For identifying live cells, LIVE/DEAD™ Fixable Violet Dead Cell Stain Kit (Invitrogen) was used. For blocking, single-cell suspensions were incubated with Mouse BD Fc Block (#553141; BD Pharmingen) in 10% FBS for 20 minutes on ice. Next, optimal concentrations of extracellular antigen fluorophore-conjugated primary antibodies were added for 20 minutes on ice (Table 1). Conjugation of antibodies was carried out using Alexa Fluor™ 488 Antibody Labeling Kit (#A20181; Invitrogen). Lastly, cells were fixed using 2% PFA. Single-cell suspension analyses were carried out on Cytek Northern Lights flow cytometer and data was processed using FlowJo software (TreeStar, Ashland, OR). During analysis, Fluorescence Minus One and isotype samples were used to determine the gating thresholds. In experiments where the treated group was compared to the control group, normalization was carried out within individual experiments to exclude inter-experimental variability. Normalization was presented as a fold-change of the control group, calculated by dividing each data point by the average of the control group for the respective experiment.

### Immunofluorescent staining of muscle and spleen

Before euthanasia, anti-CD31 antibody (see Table 1) was injected intravenously and allowed to circulate for 10 min to stain blood vessels. After cervical dislocation and removal of the gastrocnemius muscle, the muscle was fixated on a shaker using 4% PFA; 2% Triton X-100 in PBS at room temperature. Next, blocking was carried out with 5% FBS with 2% Triton X-100 in PBS on a shaker at room temperature for 48 hours. Incubation with conjugated primary antibodies in 0.2% Triton X-100 in PBS was then performed for 2 days at 4 degrees. Tissue was washed 3 x 20 min with PBS with 0.05% Tween-20 on a shaker at room temperature, followed by tissue clearing. Muscle tissue was cleared using RapiClear 1.52 (#RC152002; SUNJin Lab).

The spleen was removed and snap-frozen using liquid nitrogen. Thereafter, 20-µm-thick cryosections were prepared and fixed for 10 min in ice-cold acetone, permeabilized for 20min at room temperature with 0.1% Triton X-100 (T8787, Sigma-Aldrich), and incubated for 60 min with 10% FBS (16000044, Gibco) in DPBS (14190-094, Gibco) to block unspecific binding. After this, sections were stained with rat anti-CD169 and anti-CXCL12 antibodies for 1h at room temperature (Extended Data Table 1).

Confocal imaging was performed with an SP8 laser scanning confocal microscope (Leica). Image analysis was performed using Imaris software (Oxford Instruments).

### Human blood sampling and sample preparation

A total of 12 male patients undergoing elective repair for standard degenerative abdominal aortic aneurysm (AAA) were included (mean age, 75.8±6.5 years; range, 66.3–83.7 years). Inflammatory, infected (mycotic) and post-dissection aneurysms were excluded. Following induction of anesthesia, the abdominal aorta was exposed through a midline laparotomy and cross-clamped proximal and distal to the aneurysm. The aneurysm sac was opened and an interposition synthetic graft was sutured in place to restore blood flow ^31–33^. Arterial blood was collected into EDTA-coated tubes (#BD367525, BD Biosciences) at three time points: 1) after induction of anesthesia but before surgery, 2) between 1h and 1,5h after aortic cross-clamping, and 3) 1h after restoration of blood flow. Patient recruitment and sample collection were conducted in accordance with Ethical approval Dnr 2024-01354-01 from the Swedish Ethical Review Authority. All patients provided written informed consent. After sample collection, whole blood was mixed with HetaSep (#07906, Stemcell) and centrifuged at 200G for 5 min at room temperature, with the brake off, to isolate nucleated cells. The leukocyte-rich supernatant was harvested and any leftover red blood cells were lysed with ACK Lysis buffer (#A1049201; Gibco). For identifying live cells, LIVE/DEAD™ Fixable NIR Dead Cell Stain Kit (Invitrogen) was used. For blocking, single-cell suspensions were incubated with anti-human BD Fc Block (#564219; BD Pharmingen) in 10% FBS for 20 minutes on ice. Next, optimal concentrations of extracellular antigen fluorophore-conjugated primary antibodies were added for 20 minutes on ice (Table 1). Lastly, cells were fixed using 2% PFA. Single cell suspensions analyses were carried out on Cytek Northern Lights flow cytometer, and data was processed using FlowJo software (TreeStar, Ashland, OR). During analysis, Fluorescence Minus One and isotype samples were used to determine the gating thresholds.

## Statistical analysis

Data are presented as mean ± SEM and statistical analysis was performed using Prism 5 (GraphPad Software). Shapiro-Wilk’s test was used to assess normal distribution. Unpaired two-tailed Student’s t-test was used to compare two groups in the case of normally distributed data, and Mann-Whitney test was used if data was not normally distributed. Kruskal-Wallis with Dunn’s post-hoc test was used for comparing control data with more than two groups at one time point. Two-way ANOVA with Sidak’s post-hoc test was used to compare longitudinal measurements between two groups. Friedman test with Dunn’s multiple comparisons test was used to compare repeated measurements across three dependent related groups. P˂0.05 was considered statistically significant. Outliers were identified using Grubb’s test (Alpha=0.1), and datapoints were removed when Grubb’s test identified them as outliers for all assessed measurements. This test identified one outlier in the cohort represented in Fig. 1 and 3.

**Figure 1.**
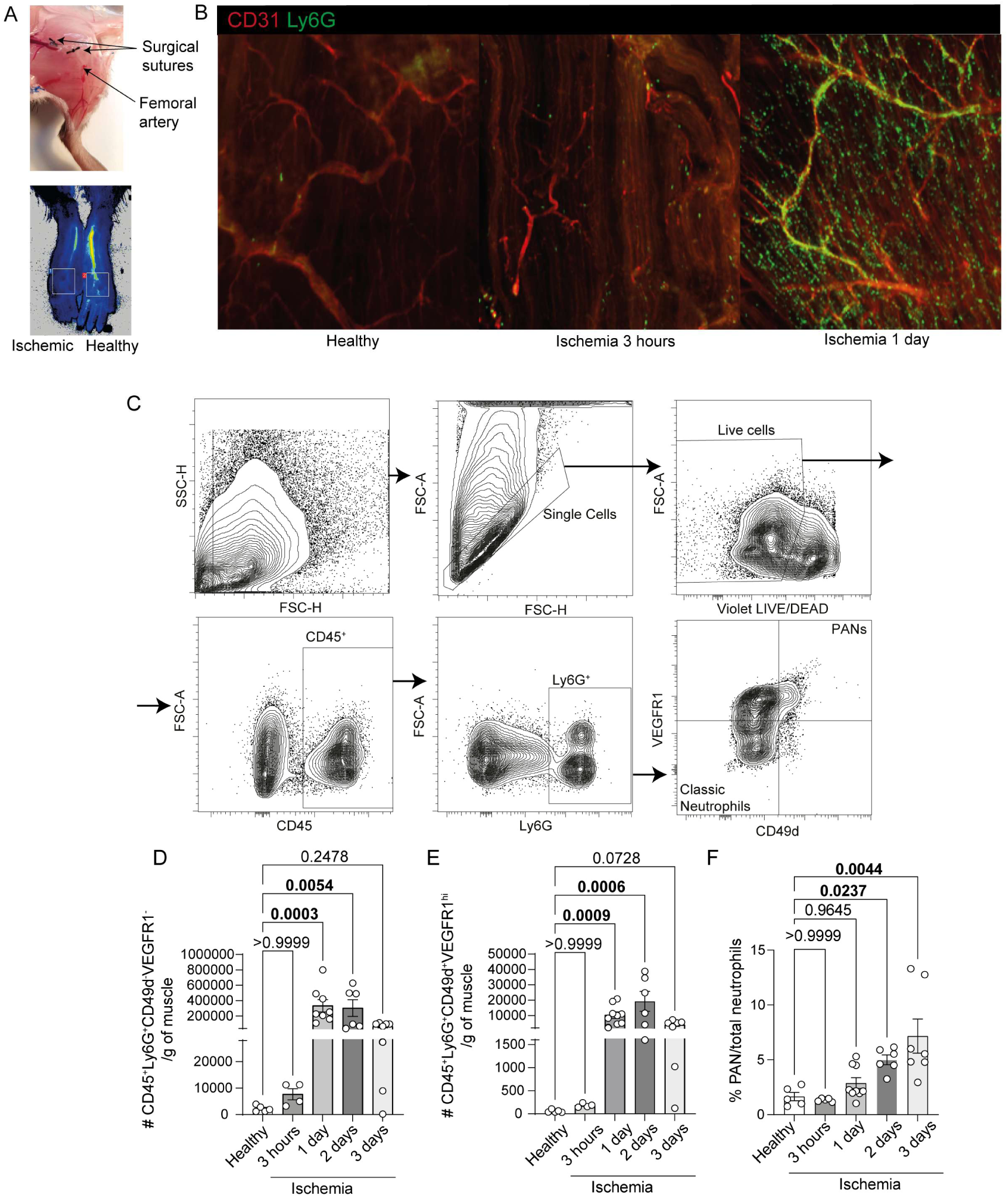
Pro-angiogenic neutrophils (PANs) are recruited to skeletal muscle following ischemia onset. A) Image of the surgical site (left) and representative blood flow of footpads after hindlimb ischemia induction measured by Laser Speckle flowmetry (right). B) Representative whole-mount images of fluorescently stained vessels (red) and neutrophils (green) of gastrocnemius muscle, healthy (left), 3 hours (center) and 1 day (right) after ischemia induction. C) Representative contour plots of gating for Ly6G^+^CD49^+^VEGFR1^hi^ (Q2) and classic Ly6G^+^CD49^-^VEGFR1^-^(Q4) neutrophils in the ischemic gastrocnemius muscle. Recruitment kinetics of D) classic neutrophils and E) Pro-angiogenic neutrophils (PANs) into the ischemic gastrocnemius muscle, along with F) proportion of PANs from total neutrophils at the different timepoints post-ischemia. Kruskal-Wallis test with Dunn’s posthoc test (D-F). Data are shown as average ± SEM.

## Results

### Pro-angiogenic Ly6G^+^CD49d^+^ VEGFR1^hi^ neutrophils are recruited to the ischemic muscle

Hindlimb ischemia (HLI) was induced by ligation of the femoral artery, which resulted in reduced blood flow to the limb downstream of the arterial ligation (Fig. 1A), and development of ischemia in e.g. the gastrocnemius muscle. While neutrophils are rare in healthy muscle, they accumulated in the ischemic gastrocnemius muscle, as evidenced by the whole-mount imaging of Ly6G^+^ neutrophils 3 hours and 1 day post-ischemia (Fig. 1B). The recruitment kinetics of CD45^+^Ly6G^+^CD49d^-^VEGFR1^-^(from here on referred to as classic neutrophils) and CD45^+^Ly6G^+^CD49d^+^VEGFR1^hi^ cells (pro-angiogenic neutrophils, PANs) to the ischemic muscle was assessed at different time points following HLI onset (3 hours, 1, 2, and 3 days) using flow cytometry (Fig. 1C-F). PANs have been previously shown to express high surface levels of CXCR4, which is internalized in response to the ligand CXCL12, particularly in dynamic disease models such as ischemia.^20,34–36^ CXCR4 levels were thus not included in the PAN gating. The numbers of both classic neutrophils and PANs per gram of muscle were significantly increased by day 1 and 2 post-ischemia (Fig. 1D-E). However, within the neutrophil population, the proportion of PAN increased over time, from 1.3 ± 0.1 % at 3 hours and 2.9 ± 0.5 % at 1 day, to 5.0 ± 1.1 % on day 2 and 7.2 ± 1.6 % on 3 days post-HLI (Fig. 1F), demonstrating a progressive enrichment of PANs during the post-ischemic response.

### Recruitment of pro-angiogenic neutrophils to the ischemic muscle is dependent on CD49d, and CD49d-inhibition results in impaired blood flow recovery of the ischemic hind limb

Pro-angiogenic neutrophils have been reported to express CD49d.^20,37^ Following CD49d-inhibition using a small molecule inhibitor given twice daily (Fig. 2A), the recruitment of classic neutrophils to the ischemic muscle was unaffected at day 3 post ischemia induction (Fig. 2B). In contrast, the accumulation of PANs was reduced by 66% in the ischemic muscle 3 days after ischemia-onset compared to the control group receiving saline (Fig. 2C). While CD49d inhibition treatment resulted in no differences in the levels of hematopoietic stem cells, endothelial progenitor cells, B cells, T cells and NK cells in the ischemic muscle (Suppl. Fig. 1A-F), it reduced the number of macrophages (CD45^+^CD11b^+^F4/80^+^, Figure 2D) as monocytes expressing CD49d are also recruited to the site of injury.^38,39^ Functional blood perfusion (heat-induced hyperemia) of the ischemic hindlimb was then assessed by laser speckle flowmetry during a topical heat challenge (Figure 2E). In saline-treated mice, the functional blood flow of the ischemic hindlimb recovered approximately 20% during the first week post-ischemia induction whereas CD49d inhibition treatment resulted in a significantly reduced recovery at 3 and 7 days after HLI (Figure 2F).

**Figure 2.**
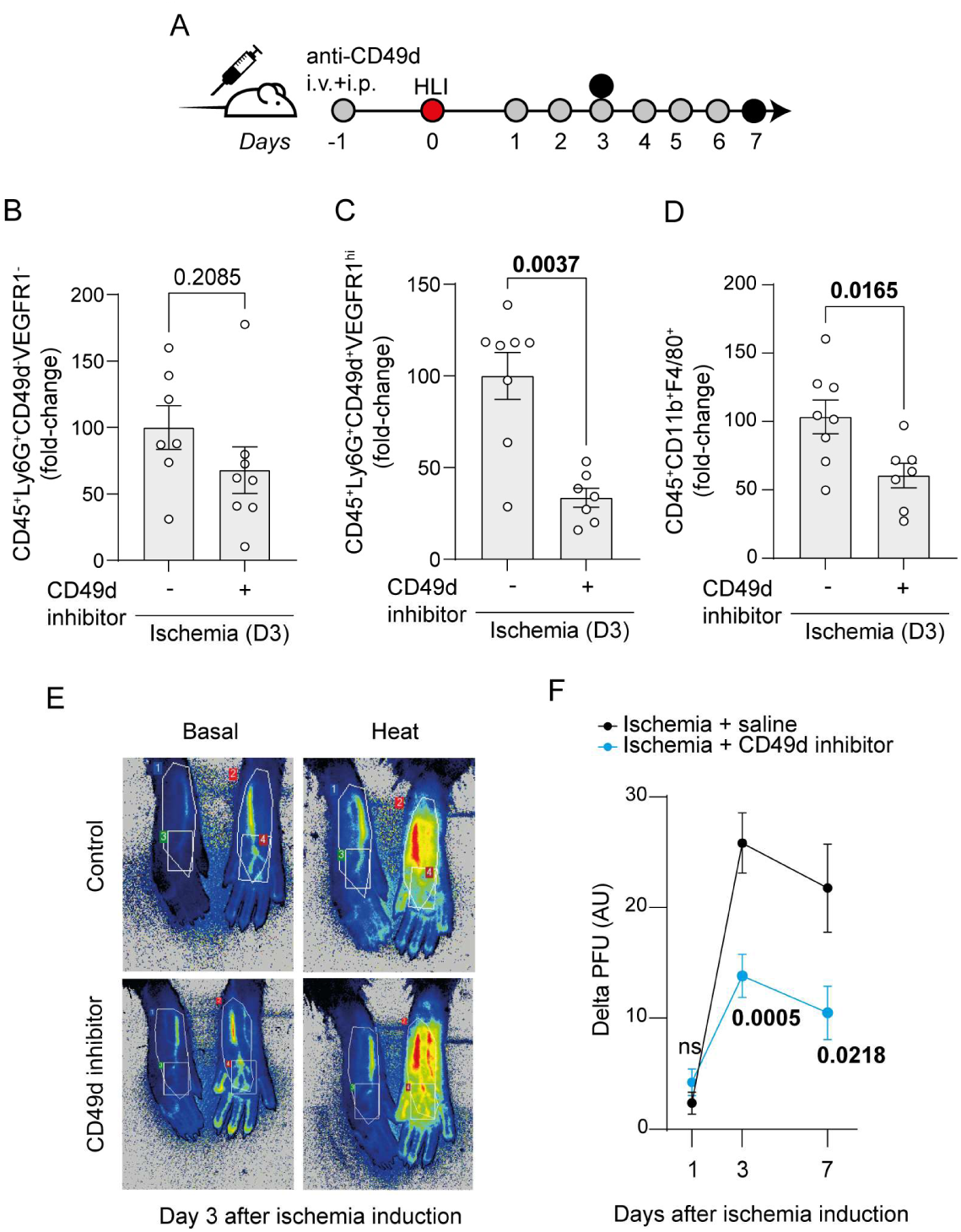
Inhibition of PAN recruitment by CD49d inhibition treatment leads to impaired blood flow recovery in the ischemic hind limb. A) Experimental layout: CD49d inhibition was achieved by intravenous (i.v.) and intraperitoneal (i.p) administration of CD49d inhibition (gray circles) one day before HLI surgery (red circle), and daily from there on for 7 days. Black circles indicate experimental endpoints. Fold changes of B) classic neutrophils, C) PANs and D) macrophages (CD45^+^CD11b^+^F4/80^+^) recruited to the ischemic gastrocnemius muscle 3 days after HLI surgery with or without treatment with CD49d inhibitors. E) Representative image of functional blood flow measurements in the hindlimb before and after heat challenge, measured by Laser Speckle flowmetry in control mice and mice treated with the CD49d inhibitor. F) Laser Speckle flowmetry measuring delta blood flow on day 1, 3, and 7 (CTRL; n=11, CD49d inhibitor; n=14) after HLI induction. Delta PFU= baseline perfusion deducted from blood flow values upon heat-challenge. Mann-Whitney U test (B), Unpaired student’s t-test (C-D), Two-way ANOVA + Sidak’s post-hoc test (F). Data are shown as average ± SEM.

**Figure 3.**
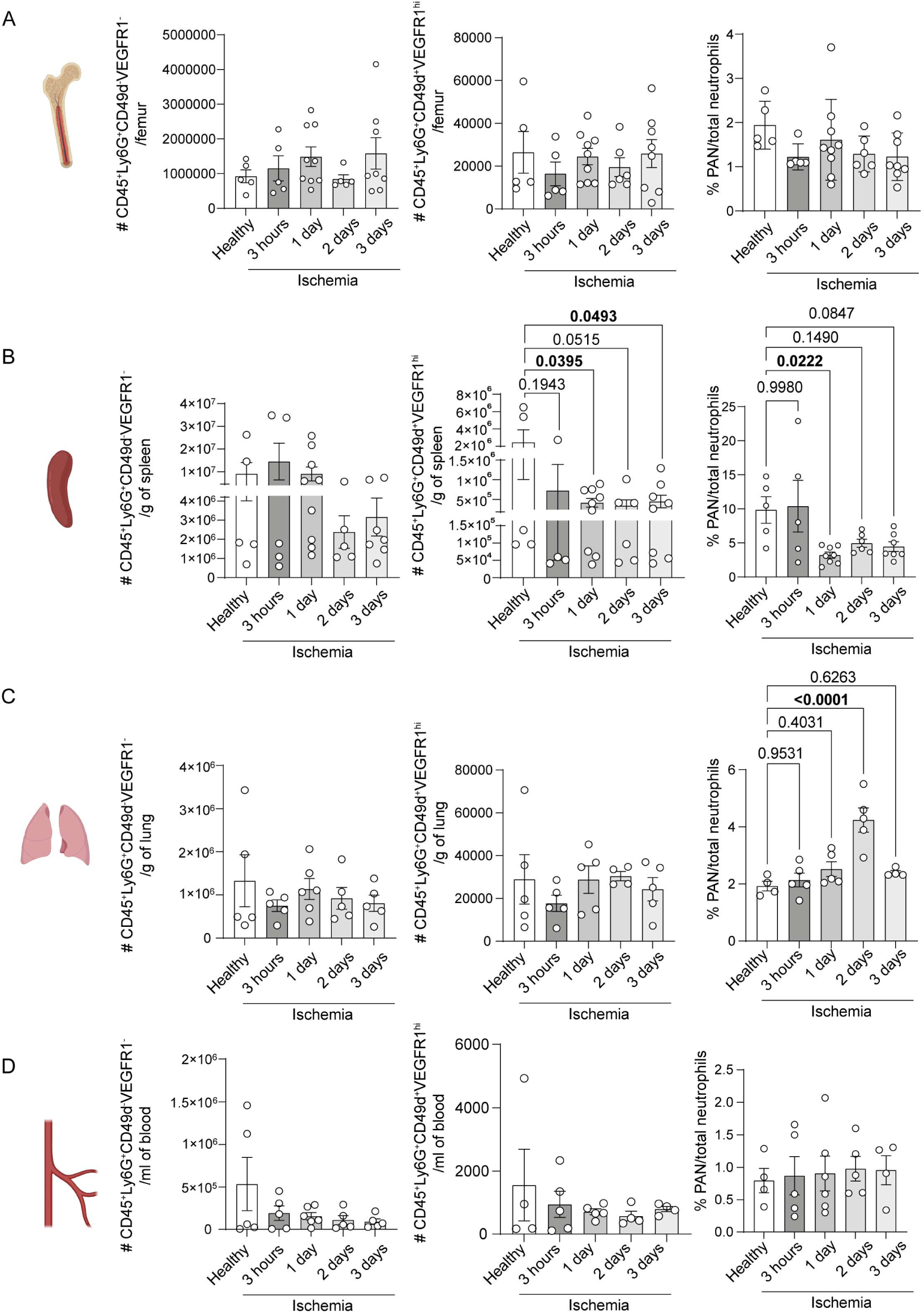
Steady-state levels of classic neutrophils and PANs in the bone marrow, spleen, lungs and circulation, and their kinetics upon HLI induction reveal the spleen as a peripheral reservoir of PANs. A) Kinetics of classic CD49d^-^VEGFR1^-^neutrophils and CD49d^+^VEGFR1^hi^ PANs in A) bone marrow, B) spleen, C) lungs and D) circulation. Kruskal-Wallis test with Dunn’s post-hoc test (A, D), Ordinary one-way ANOVA with Dunnett’s post-hoc test (B-C). Data are shown as average ± SEM.

### Pro-angiogenic neutrophils are mobilized from the spleen at ischemia onset

To investigate if the neutrophil subsets are stored in peripheral reservoirs at steady state^12,24,40^, classic neutrophils and PANs from bone marrow (Fig. 3A, Suppl. Fig. 2A), spleen (Fig. 3B, Suppl. Fig. 2B), lungs (Fig. 3C, Suppl. Fig. 2C) and circulation were evaluated (Fig. 3D, Suppl. Fig. 2D). Interestingly, high numbers of PANs, as well as their proportion of the total neutrophil population, were identified in the spleen (9.8±1.9%), in contrast to the other investigated organs (bone marrow: 1.9±0.2%, lungs: 1.9±0.2%, circulation: 0.8±0.2%, Fig. 3A-D). Upon ischemia induction, the numbers of PANs were significantly reduced in the spleen at 1 and 3 days post-ischemia, (Fig. 3B), whereas the levels of classic neutrophils remained unaltered. As a result, the proportion of PANs out of total neutrophils was reduced at 1 and 3 days post-ischemia onset (3.2±0.4% and 4.4±0.7% of total neutrophils, at days 1 and 3, respectively). Furthermore, the levels of classic neutrophils or PANs did not change in bone marrow or circulation in response to ischemia. Interestingly, the proportion of PANs was increased in the lung 2 days post-HLI (4.2±0.4% of total neutrophils, Fig. 3C).

To assess the importance of the spleen as a source of PANs following ischemic insults, splenectomy was performed on mice seven days prior to HLI induction (experimental layout, Fig. 4A). One day following ischemia onset, splenectomised mice had similar levels of classic neutrophils in their ischemic muscle when compared to mice that were sham-splenectomised (Fig. 4B). In contrast, the number of PANs was significantly reduced by more than 50% in the ischemic gastrocnemius muscle of splenectomised mice when compared to sham group (Fig. 4C) even though still increased when compared to mice that received sham-HLI with or without splenectomy (Fig. 4D), demonstrating that a significant proportion of these cells are mobilized from the spleen in response to ischemia onset. In parallel, reduced PAN proportion was observed in the bone marrow of splenectomised but not in sham-splenectomised mice at 1 d post-ischemia induction, indicating that the bone marrow compensates for the lack of spleen (Fig. 4E). Together, these results demonstrate that the spleen serves as a peripheral pool of PANs, which are promptly released to circulation upon ischemia induction for the recruitment into the injured tissue.

**Figure 4.**
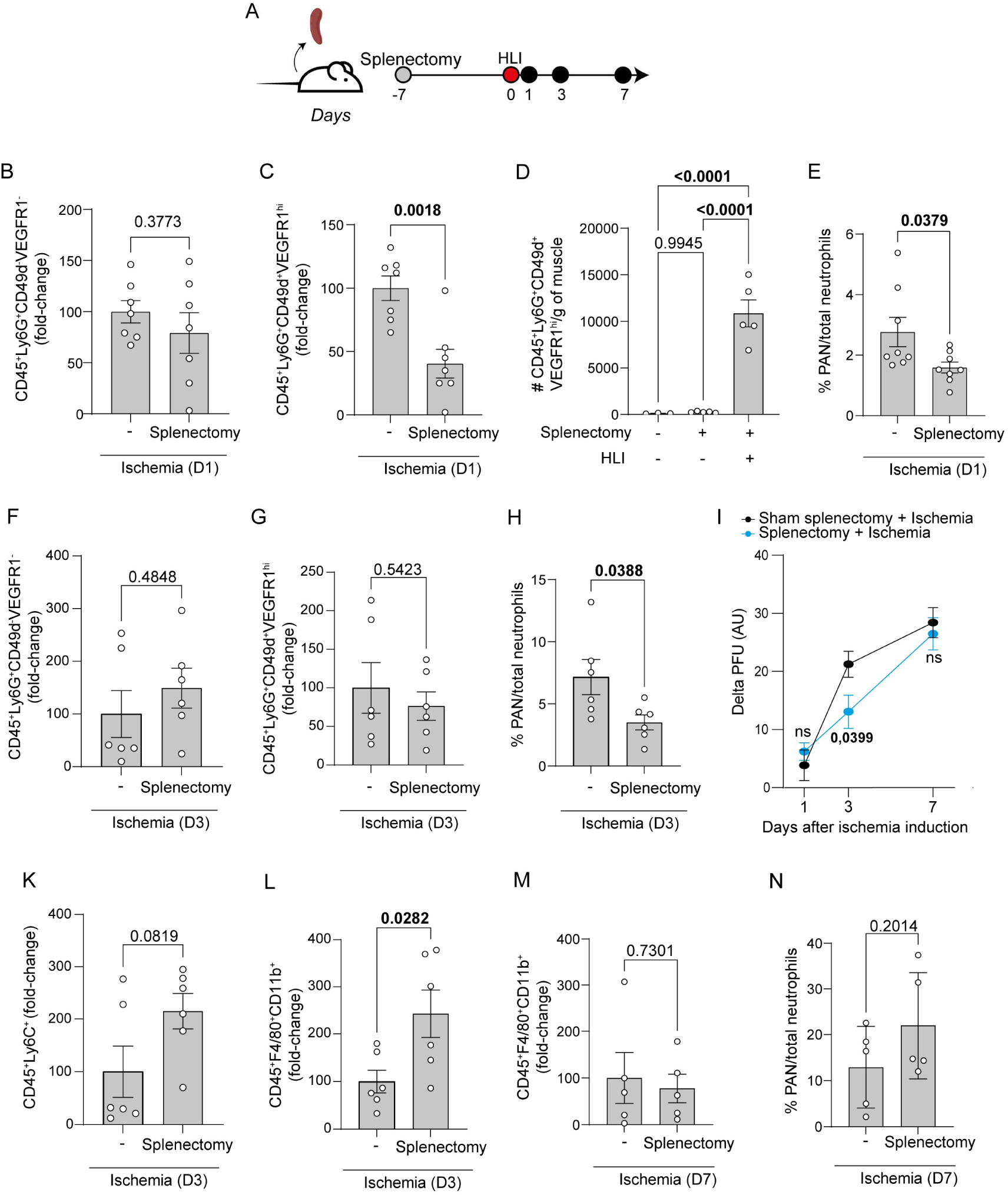
PANs recruited to ischemic muscle are mobilized from the spleen upon ischemia onset. A) Experimental layout: splenectomy (grey circle) was carried out 7 days prior to HLI induction (red circle). Black circle indicates laser speckle measurements and experimental endpoints at days 1, 3 and 7. Fold changes of B) classic neutrophils and C) PANs recruited to the ischemic muscle of splenectomised and sham-operated mice 1 day post-HLI. D) Number of PANs in controls (sham-splenectomy, sham-HLI and splenectomised, sham-HLI), and splenectomised ischemic mice 1 day post-HLI. E) Proportion of PANs in the bone marrow of control and splenectomised mice 1 day post-HLI. Fold changes of F) classic neutrophils and G) PANs recruited to the ischemic muscle of splenectomised and sham-operated mice 3 days post-HLI. H) Proportion of PANs in the ischemic muscle of control and splenectomised mice 3 days post-HLI. I) Laser Speckle flowmetry measuring delta blood flow on day 1, 3, and 7 (sham; n=6, splenectomy; n=9) after HLI induction. Delta PFU= baseline perfusion deducted from perfusion after blood flow regulation upon heat challenge. Fold changes of K) monocytes and L) macrophages recruited to the ischemic muscle of splenectomised and sham-operated mice 3 days post-HLI. M) Fold changes of macrophages recruited to the ischemic muscle of splenectomised and sham-operated mice 7 days post-HLI. N) Proportion of PANs in the ischemic muscle of control and splenectomised mice 7 days post-HLI. Unpaired student’s t-test (B-C, G-H, K-N). Ordinary one-way ANOVA with Dunnett’s post-hoc test (D). Mann-Whitney U test (E-F). Two-way ANOVA + Sidak’s post-hoc test (I). Data are shown as average ± SEM.

Three days following ischemia induction, the splenectomised mice had similar levels of both classic neutrophils and PANs in their ischemic muscle (Fig. 4F-G), whereas the proportion of PANs was still significantly decreased in the ischemic muscle (Fig. 4H). Moreover, laser speckle flowmetry revealed reduced functional blood perfusion (heat-induced hyperemia) in splenectomised mice compared to the control group at 3 days after HLI (Figure 4I). The macrophages (CD45^+^F4/80^+^CD11b^+^), but not number of monocytes (CD45^+^Ly6C^+^), was increased in the injured gastrocnemius in splenectomised mice when compared to sham-operated mice (Fig. 4K-L). This observation is opposite to what was observed following CD49d inhibition treatment (Fig. 2D), and suggests that the impaired blood flow restoration following splenectomy is a macrophage-independent effect and due to the reduced recruitment of PANs (Fig. 4H). By 7 days post-HLI, functional blood perfusion no longer differed between the two groups (Fig. 4I), which coincided with that PANs and macrophage levels in the injured muscle of splenectomised mice were no longer different from those in sham-operated mice (Fig. 4M– N). Overall, these experiments reveal the spleen as source of pro-angiogenic neutrophils that are mobilized to the ischemic muscle to improve blood flow recovery.

### Mobilization of pro-angiogenic neutrophils from the spleen is regulated by sympathetic signaling-dependent reduction of CXCL12 expression in the splenic red pulp

The spleen is innervated by sympathetic neurons which regulate immune cell localization, activation, and effector functions by noradrenergic signaling, thereby providing a key mechanism through which the nervous system modulates splenic innate and adaptive immunity.^41–43^ To investigate the involvement of sympathetic neurons in the splenic egress of PANs, systemic sympathectomy was achieved by injections of 6-OHDA, a catecholaminergic neuron-selective toxin, every second day during the week prior to ischemia induction (Fig. 5A). Consequently, the effect of systemic sympathectomy on the recruitment of neutrophils to the ischemic muscle was investigated. Whereas no significant effect was observed for classic neutrophils (Fig. 5B), the levels of PANs were significantly reduced within the ischemic muscle 1 day post-HLI (Fig. 5C) in systemically sympathectomized mice. In parallel, systemic sympathectomy did not affect splenic levels of classic neutrophils following HLI induction (Fig. 5D), but selectively increased splenic PAN levels by 2.8-fold at 1 day post-HLI (Fig. 5E). In addition, while the bone marrow of 6-OHDA-treated mice did not differ in the number of classic neutrophils, the numbers of PANs were reduced post-ischemia induction (Suppl. Fig. 3A-C) similar to the observations in splenectomised mice (Fig. 4E), demonstrating ischemia-induced mobilization of PANs from bone marrow. These observations translated into impaired blood flow recovery at day 1 post-ischemia in sympathectomized mice compared to the control group but not at later time points (Fig. 5F).

**Figure 5.**
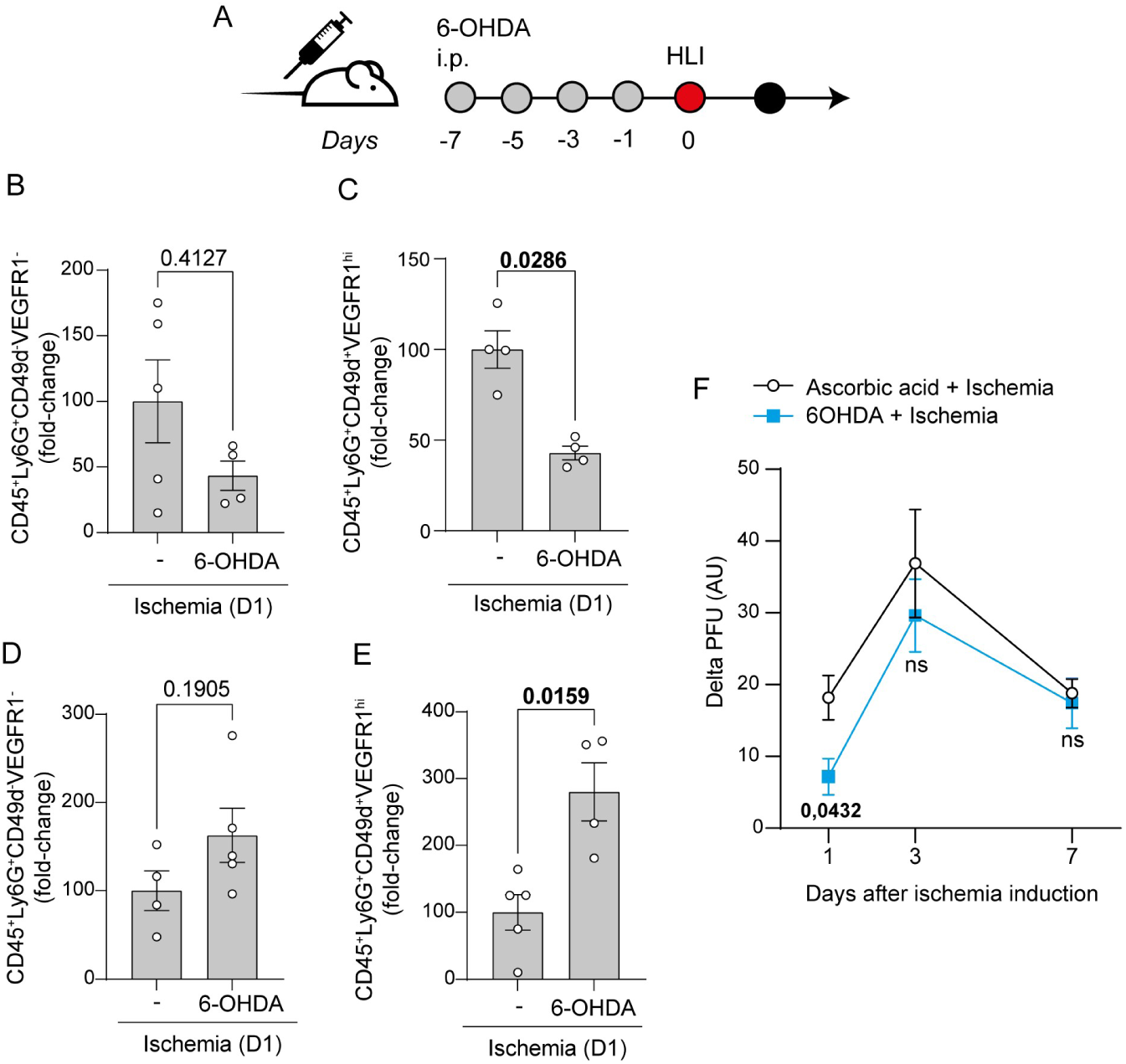
Mobilization of pro-angiogenic neutrophils from the spleen is regulated by sympathetic signaling. A) Experimental layout: systemic sympathectomy was achieved by intraperitoneal 6-OHDA injection (grey circles) every 48 hours starting 7 days before HLI surgery (red circle). Black circle indicates experimental endpoint at day 1 post-HLI. Normalised percentages of B) classic neutrophils and H) PANs in the ischemic muscle of systemically sympathectomised (6-OHDA) and control treated mice. Normalised percentages of D) classic neutrophils and E) PANs in the spleens of systemically sympathectomised (6-OHDA) and control treated mice 1 day post-HLI. F) Laser Speckle flowmetry measuring delta blood flow on day 1, 3, and 7 (control; n=11, 6-OHDA; n=12) after HLI induction. Delta PFU= baseline perfusion deducted from perfusion after blood flow regulation upon heat-challenge. Mann-Whitney U test (B-E). Two-way ANOVA + Sidak’s post-hoc test (F). Data are shown as average ± SEM. P-values are displayed in the graphs.

To isolate the effects of the sympathetic signaling to the spleen, splenic denervation was performed on mice 7 days prior to HLI induction (experimental layout, Fig. 6A). Similar to the observations made in systemically sympathectomised mice (Fig. 5B-C) and splenectomised mice (Fig. 4B-C), spleen-specific denervation did not affect the recruitment of classic neutrophils to the ischemic muscle 1 day post-ischemia induction (Fig. 6B), whereas the recruitment of PANs was reduced (Fig. 6C). Moreover, at day 3 post-ischemia induction the number of classic neutrophils was increased in mice that had undergone splenic denervation (Fig. 6D), whereas the number of PANs was not altered (Fig. 6E), resulting in that the proportion of PANs within the ischemic muscle was reduced 3 days post-HLI (Fig. 6F). Alongside the increase in classic neutrophils in the ischemic muscle, the number of monocytes but not macrophages also increased at day 3, consistent with an ongoing inflammatory response and active recruitment of circulating myeloid cells to the injured tissue (Fig. 6G-H).^44^ The blood flow recovery to the hindlimb in splenically denervated mice was impaired, as functional blood perfusion (heat-induced hyperemia) assessed by laser speckle flowmetry was reduced when compared to the control group 3 and 7 days after HLI (Figure 6I), again demonstrating the importance of PANs in the recovery from injury.

**Figure 6.**
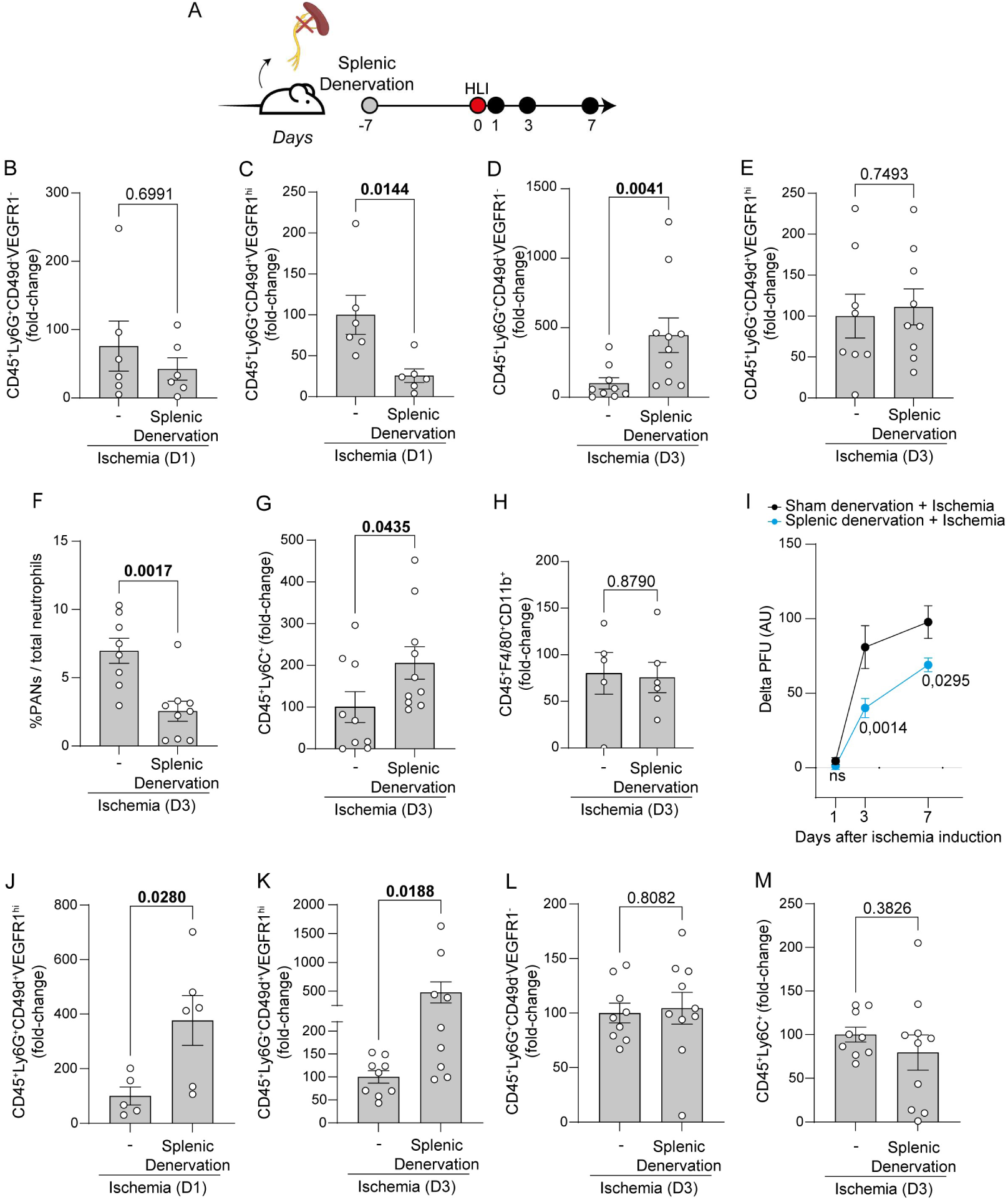
Splenic denervation impairs the recruitment of pro-angiogenic neutrophils into the ischemic muscle and induces splenic retention of these cells. A) Experimental layout: surgical splenic denervation was performed (grey circles) 7 days before HLI surgery (red circle). Black circle indicates both laser speckle measurements and experimental endpoints post-HLI. Fold changes of B) classic neutrophils and C) PANs in the ischemic muscle of sham-operated and splenic denervated mice 1 and 3 days (D and E, respectively) post-HLI. F) Percentages of PANs in the ischemic muscle 3 days post-ischemia. Fold-change of G) monocytes and H) macrophages in the ischemic muscle of splenic denervated and sham-operated mice 3 days post-HLI. I) Laser Speckle flowmetry measuring delta blood flow on day 1, 3, and 7 (sham; n=6, splenic denervation; n=9) after HLI induction. Delta PFU= baseline perfusion deducted from blood flow upon heat-challenge. Fold changes of PANs in the spleen of splenic denervated and sham-operated mice 1 day (J) and 3 days (K) post-HLI. Fold changes of L) classic neutrophils and M) monocytes in the spleen of splenic denervated and sham-operated mice 3 days post-HLI. Unpaired student’s t-test (C, E-F, J-K). Mann-Whitney U test (B, D, G-H, L-M). Two-way ANOVA + Sidak’s post-hoc test (I). Data are shown as average ± SEM.

The neutrophils in the spleen were then assessed in the mice undergoing both splenic denervation and induction of ischemic hindlimb. Both at 1 and 3 days post-HLI, the number of PANs in the spleen of denervated mice was increased when compared to mice with intact splenic innervation (Fig. 6J-K), whereas the levels of classic neutrophils and monocytes were not altered (Fig 6L-M). These results demonstrate that splenic innervation mediates the mobilization of PANs but not classic neutrophils or monocytes from the spleen, which influence the concomitant PAN recruitment to and blood flow restoration of sites of ischemia.

In contrast to mature classic neutrophils in circulation, circulating PANs have been reported to express high levels of CXCR4^20^, a chemokine receptor for the chemokine CXCL12. CXCL12 is a retention factor regulating the mobilization of mature neutrophils from the bone marrow hematopoietic compartment into the circulation.^45^ To investigate whether splenic expression of CXCL12 retains PANs in the spleen, and if PAN mobilization from spleen in response to peripheral ischemia is regulated by CXCL12, immunohistochemistry was performed on spleens isolated from control mice and mice following ischemia induction (Fig. 7A). Using CD169^+^ metallophilic marginal zone macrophages to identify the border between the red and the white pulp^46^, we found that CXCL12 was expressed in the red pulp where neutrophils and macrophages are known to reside (3.2±0.5 % area), but were not detected in the white pulp (composed of mainly B and T cells, Fig. 7B) during homeostasis. Upon induction of ischemia, the levels of red pulp CXCL12 were significantly reduced (1.0±0.4 % 3 hours post-HLI, Fig. 7C). Together, these results identify sympathetic innervation as an important regulator of splenic PAN mobilization and suggest that ischemia-induced regulation of CXCL12 in the splenic red pulp contributes to their release.

**Figure 7.**
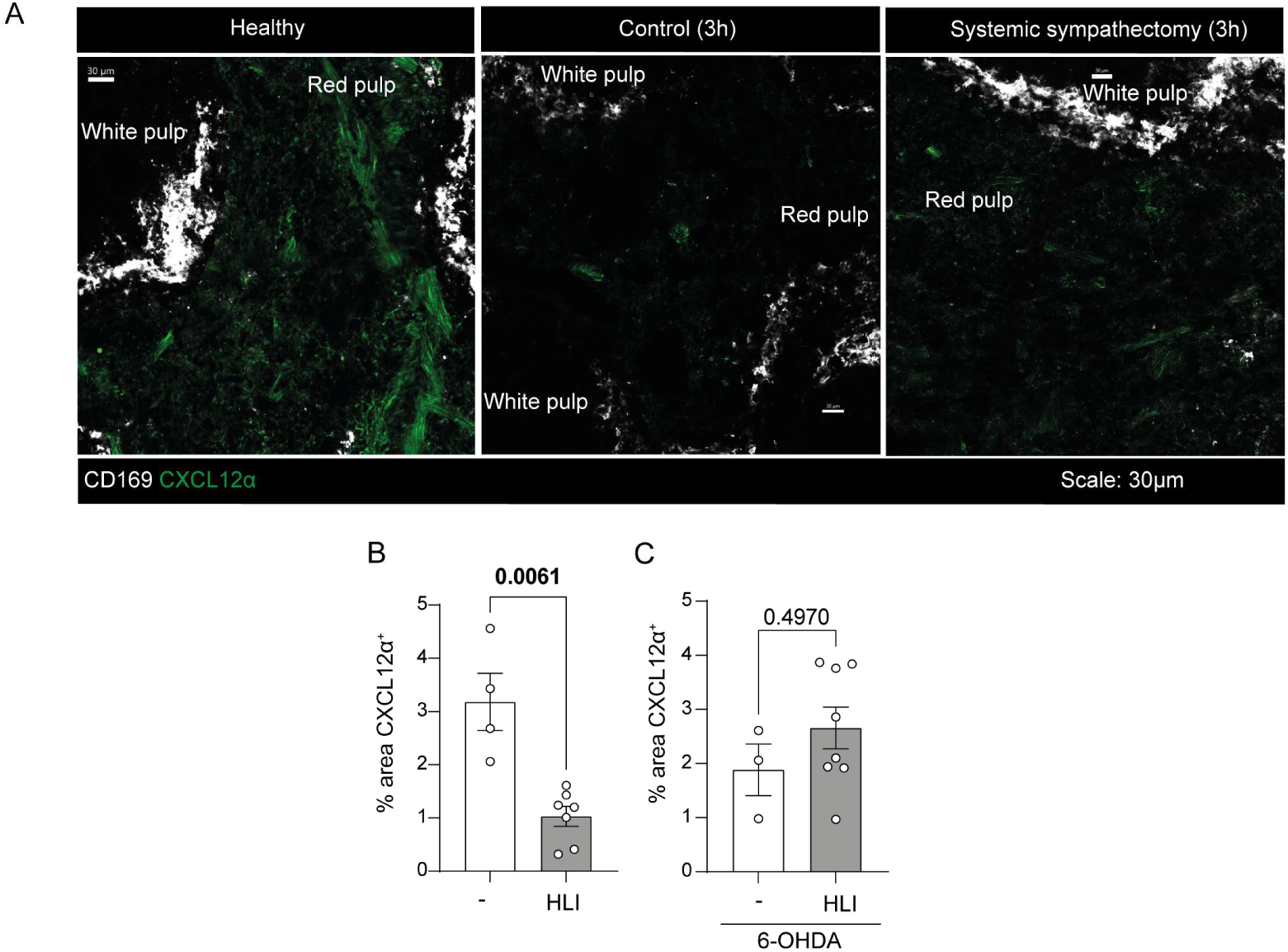
Mobilization of pro-angiogenic neutrophils from the spleen is regulated by sympathetic signaling-dependent reduction of CXCL12-expression in the splenic red pulp. A) Representative images of CD169 (white) and CXCL12 (green) immunohistochemical staining in the spleen at basal state, 3 hours after ischemia in control-treated mice and 3 hours after ischemia in systemically sympathectomised mice. Scale bar: 30 µm. B) Percentage of area CXCL12^+^ in the red pulp of control-treated mice at 3 hours post-HLI. C) Percentage of area CXCL12^+^ in the red pulp of 6-OHDA-treated (right) mice at 3 hours post-HLI. Mann-Whitney U test (B-C). Data are shown as average ± SEM. P-values are displayed in the graphs.

### Mobilization of pro-angiogenic neutrophils to circulation in humans is stimulated by ischemia

To investigate whether our observations in mice could be translated to humans, circulating pro-angiogenic neutrophils, previously identified as CD45^+^CD14^-^CD16^+^CD11b^+^VEGFR1^+^ CD49d^+^^20^ (Suppl. Fig. 4A) were assessed in patients undergoing elective AAA repair, during which the aorta is temporarily cross-clamped, resulting in acute lower-body ischemia. Pro-angiogenic neutrophils were isolated from blood samples by gradient centrifugation and quantified by flow cytometry following collection before surgery, 1 – 1.5 hours after aortic cross-clamping, and 1 hour after restoration of blood flow. The sampling period spanned approximately 5 h, from the beginning to the end of surgery (Fig. 8A).

**Figure 8.**
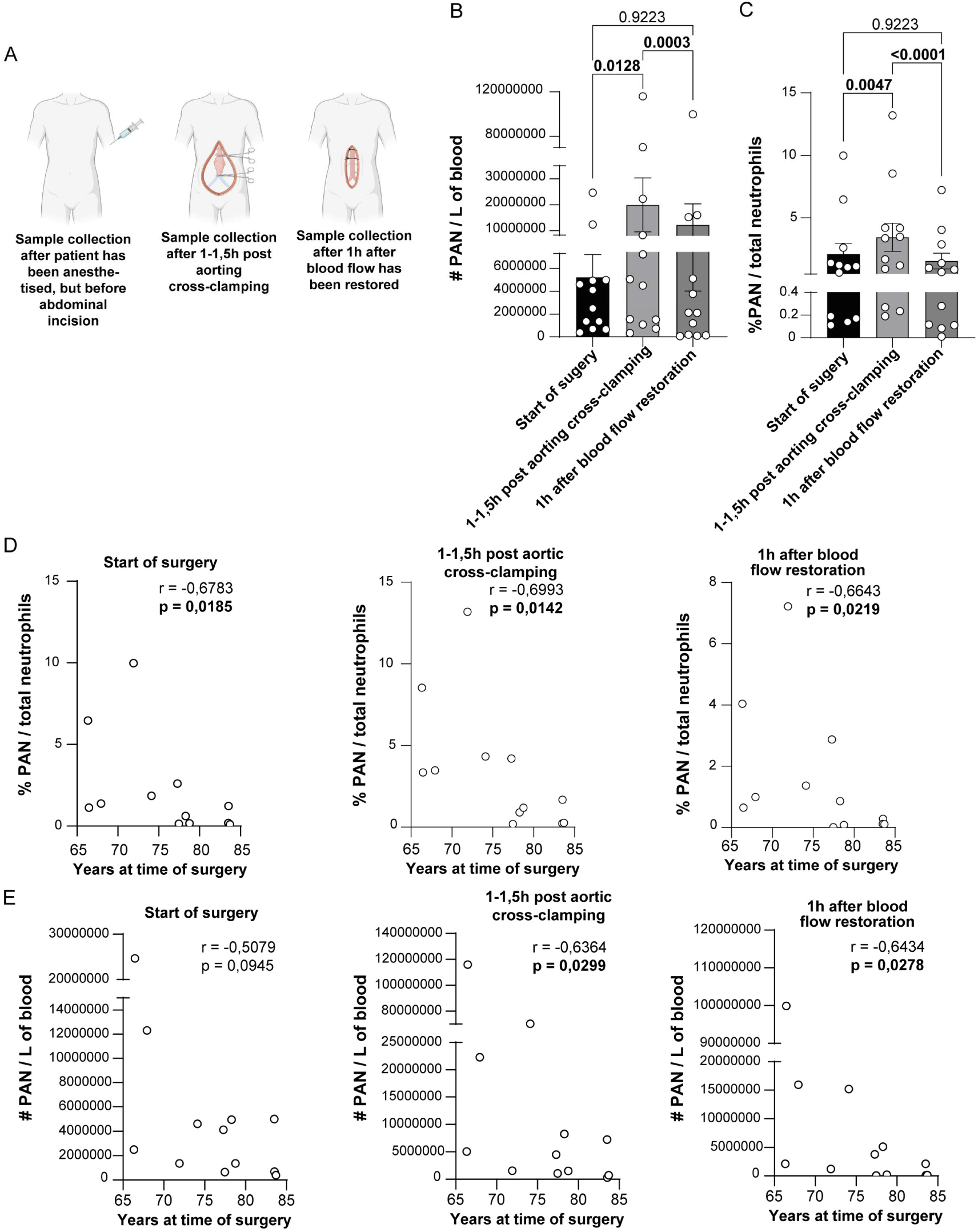
Aorta ligation induced the egress of pro-angiogenic neutrophils into circulation. A) Experimental layout for sample collection during abdominal aortic aneurysm repair. B) Numbers of PAN per liter of blood through surgical timepoints. C) Percentage of PAN from total neutrophils through surgical timepoints. D) Correlation between age and percentage of PAN from total neutrophils. E) Correlation between age and numbers of PAN per litre of blood. Friedman test with Dunn’s multiple comparisons test (B-C). Data are shown as average ± SEM. Nonparametric Spearman correlation test (D-E).

For the 12 male patients included in this study (mean age, 75.8±6.5 years; range, 66.3–83.7 years), the neutrophil population constituted 56.7±5.7% of the immune cells present in the blood at the start of surgery (Suppl. Fig. 4B), At this time point, the PAN levels were 2,1±0,9%, which is within the previously described range (numbers: 5.2×10^6^±2.0×10^6^ cells/L, Fig. 8B-C).^20^

After 1.5 h of aortic cross-clamping, PAN levels increased significantly by 3,8-fold (2.0×10^7^±1.0×10^7^ cells/L), as the proportion of PANs which increased to 3.5±1.1% (Fig. 8B-C). One hour after restoration of blood flow, the PAN levels had decreased significantly compared with the cross-clamping time point (1.2×10^7^±8.1×10^6^ cells/L), and the proportion of PANs decreased to 1,6% (Fig. 8B-C). Furthermore, the total neutrophil proportions did not change throughout the studied time points (Suppl. Fig. 4B). These findings demonstrate that, consistent with our observations in mice, PANs are rapidly mobilized in response to acute ischemia.

Furthermore, as both bone marrow and spleen undergo substantial age-related changes that might affect immune cell homeostasis, we investigated whether patient age influenced the abundance and ischemia-induced mobilization of circulating PANs.^47,48^ Overall, increasing age was associated with lower circulating PAN levels, with the proportion of PANs being lower in patients aged >77 years than in younger patients (Spearman correlation test, Fig. 8D-E). Although PAN numbers did not correlate with age at the start of surgery, significant inverse correlations emerged after aortic cross-clamping and persisted after restoration of blood flow (Fig. 8E). Similarly, the proportion of PANs was inversely correlated with patient age at all three time points (Fig. 8D). To understand if this age-related decrease could be motivated by a general decrease in neutrophils, the percentage of neutrophils and PAN from total immune cells were correlated to patient age. Whilst the percentage of total neutrophils did not correlate with patient age (Suppl. Fig. 4C), the percentage of PAN from total immune cells remains strongly correlated to age (Suppl. Fig. 4D). Collectively, these findings suggest that the capacity to mobilize PANs in response to acute ischemia declines with age, potentially contributing to impaired vascular repair in older individuals.

## Discussion

Our findings reveal a previously unrecognized role for the spleen in blood flow recovery during ischemia, serving as a major reservoir of pro-angiogenic neutrophils (PANs) that are rapidly mobilized in response to injury. We demonstrate that PANs accumulate in ischemic skeletal muscle in a CD49d-dependent manner and contribute to restoration of limb perfusion. Furthermore, we identify that sympathetic regulation of splenic CXCL12 directs PAN release from the splenic red pulp, providing a mechanistic link between neural activation following ischemia and mobilization of reparative immune cells. Finally, the rapid mobilization of a corresponding PAN population in human blood following acute lower-limb ischemia further supports the translational relevance of this response.

Previous studies have identified PANs as a specialized subset of circulating neutrophils in mice and humans, characterized by expression of CD49d, VEGFR1 and CXCR4.^18,20^ In experimental mouse models, PANs have been shown to promote angiogenesis through the release of matrix metalloproteinase-9 following VEGF-A-dependent recruitment, while inhibition of their recruitment impaired vascularization of transplanted pancreatic islets.^18,20^ Here, we extend these findings to hindlimb ischemia, a mouse model of peripheral artery disease, and demonstrate that PANs rapidly accumulate in ischemic skeletal muscle and contribute to restoration of tissue perfusion.

A central finding of our study is the identification of the spleen as a major extramedullary reservoir of PANs. While neutrophils are primarily produced in the bone marrow, increasing evidence suggests that extramedullary neutrophil pools are maintained in organs such as the spleen and lungs.^12,24,40,49^ We found that PANs were particularly enriched in the spleen, where they constitute a substantially larger fraction of the neutrophil population than in bone marrow, lungs, or circulation. Following ischemia, splenic PAN numbers rapidly declined, coinciding with their accumulation in ischemic muscle. Importantly, splenectomy markedly reduced PAN recruitment to ischemic muscle and impaired early blood flow recovery, demonstrating the functional importance of this reservoir. These observations occurred despite increased macrophage recruitment to the site of ischemia, further supporting a contribution of PANs to vascular recovery that cannot readily be compensated for by other inflammatory cell populations. Nevertheless, PANs were still detected in the ischemic tissue of splenectomised mice, indicating that the spleen is a major but not exclusive source. In agreement with previous reports of compensatory myeloid cell for the loss of splenic myeloid reservoirs during inflammatory responses^24^, we observed reduced PAN levels within the bone marrow following ischemia in splenectomised mice, consistent with enhanced mobilization from this compartment. However, this compensatory response was insufficient to restore normal perfusion recovery. Whether this reflects inadequate numbers of mobilized PANs, their mobilization kinetics, or their functional properties remains unknown.

Beyond its role as a reservoir, the spleen may also provide an environment that contribute to the maintenance or functional programming of PANs. Previous work has demonstrated that neutrophils acquire tissue-specific transcriptional programs following residency within peripheral organs.^50^ This possibility is particularly intriguing given the relative low oxygen tension of the splenic red pulp^51^, as hypoxia has been shown to induce expression of both CXCR4 and VEGFR1 in multiple cell types, including monocytes, macrophages, endothelial cells, and mesenchymal stromal cells.^52,53^ However, our findings do not establish whether PANs acquire their pro-angiogenic phenotype within the spleen or are generated elsewhere and selectively retained. Defining the transcriptional and functional properties of PANs from different anatomical reservoirs will be necessary to determine whether they are transcriptomically and functionally equivalent.

Our findings further identify sympathetic signaling as an important regulator of PAN release from the splenic reservoir. The spleen is densely innervated by sympathetic nerve fibers, and neural regulation of immune cell trafficking has previously been demonstrated in several settings including of antibacterial immunity in the spleen, circadian leukocyte trafficking, neutrophil mobilization from bone marrow, and splenic monocyte release following ischemic stroke.^42,43,54–56^ Here, we observed rapid reduction of CXCL12 expression within the splenic red pulp following ischemia. The CXCL12-CXCR4 axis is a well-established regulator of neutrophil retention in the bone marrow, where reduced CXCL12 signaling promotes neutrophil egress into the circulation.^45,57–59^ Importantly, systemic sympathectomy prevented ischemia-induced downregulation of splenic CXCL12 and was associated with PAN retention within the spleen and reduced recruitment to ischemic tissue. Selective splenic denervation similarly increased splenic PAN abundance, reduced their accumulation in ischemic muscle, and impaired blood flow recovery. Together, these findings suggest that sympathetic signals regulate PAN mobilization through modulation of splenic CXCL12 levels, indicating that retention mechanisms previously characterized in the bone marrow may also operate within the spleen. Notably, disruption of splenic sympathetic signaling impaired blood flow recovery despite increased recruitment of monocytes and macrophages, further supporting a distinct contribution of PANs to the restorative response following ischemic injury. The cellular and molecular mechanisms connecting sympathetic activation to reduced levels of splenic CXCL12 remain to be defined. CXCL12 is known to be produced by multiple stromal cell populations within the spleen, including fibroblastic reticular cells and vascular stromal cells.^60,61^ Moreover, β-adrenergic receptor signaling has been implicated in the regulation of CXCL12 expression in hematopoietic niches, and similar mechanisms may therefore contribute also to ischemia-induced remodeling of splenic retention signals.^57,60^

To assess the relevance of our findings to humans, we examined circulating PANs in patients undergoing temporary aortic cross-clamping during abdominal aortic aneurysm repair. We identified a corresponding neutrophil population, characterized by expression of CD49d and VEGFR1, that increased rapidly following induction of ischemia and declined following reperfusion. These findings indicate that acute ischemia is associated with rapid changes in circulating PAN numbers in humans, consistent with mobilization of a pre-existing population across species, even though neither the anatomical source of these cells nor their contribution to vascular repair in humans has been established. However, previous studies have reported long-term associations between splenectomy and cardiovascular complications, including acute coronary syndrome and thrombosis^62–64^, consistent with a broader role for the spleen in cardiovascular homeostasis. Whether the human spleen represents a comparable reservoir, and whether similar neuro-immune mechanisms regulate PAN mobilization will require further investigations.

The regulated mobilization of specialized neutrophil populations may represent a broader feature of the systemic response to ischemic injury. A recent study of patients with ST-elevation myocardial infarction identified mobilized immature neutrophil progenitors expressing both CXCR4 and CD49d, and that the magnitude of mobilization of this specific neutrophil subset was positively associated with disease severity and inflammatory activation.^48^ Although these cells differ from the mature PAN population investigated here due to their low expression of CD16, the findings suggest that ischemic injury can induce distinct neutrophil mobilization responses. Whether these populations share regulatory pathways or developmental relationships remain to be established.

We additionally observed an age-associated reduction in circulating PAN abundance and ischemia-induced PAN mobilization. Given that aging is accompanied by substantial alterations in splenic architecture and immune function^47^, these observations raise the possibility that age-related changes in neutrophil reservoirs or mobilization pathways influence the response to ischemic injury. Further studies will be required to determine whether impaired PAN mobilization contributes to age-related differences in recovery following ischemic events.

Our findings suggest that modulation of PAN mobilization or recruitment could provide therapeutic opportunities to enhance vascular repair. The observation that PAN mobilization is regulated by sympathetic signaling and that PAN accumulation in ischemic tissue supports vascular recovery suggests that therapeutic modulation of PAN mobilization, recruitment, or function could represent a novel strategy to enhance tissue repair. Whereas current revascularization strategies primarily focus on reopening occluded arteries and restoring macroscopic blood flow, approaches targeting reparative immune cell populations may complement these treatments by supporting microvascular recovery ^48,65^. This concept is consistent with our previous work demonstrating that distinct myeloid cell populations can promote restored functional blood perfusion in ischemic tissues ^28,66^. We previously identified a population of macrophages that accumulate perivascularly at sites of ischemia, where they induce transdifferentiation towards pericytes and enhance vascular maturation and perfusion recovery following ischemic injury.^28,66^ Complementary, recent studies have identified proangiogenic monocytes (PAMs) as a reparative myeloid population with potent vascular regenerative properties.^67^ Building on these observations, we and others have demonstrated that therapeutically targeting reparative myeloid cell programs can enhance revascularization and tissue recovery following ischemic injury.^67,28^ Together, these studies and the present findings support a broader paradigm in which selective mobilization or activation of endogenous reparative myeloid subsets can improve recovery after ischemic injury.

Importantly, our current work extends these observations by identifying the sympathetic spleen-PAN axis as a mechanism regulating the availability of reparative immune cells after ischemia. Although PANs and restorative macrophages differ in lineage and effector functions, both populations appear to support tissue repair and vascular regeneration, suggesting that targeted modulation of endogenous reparative myeloid cell subsets may represent a promising therapeutic strategy. However, given the pro-angiogenic properties of both PANs and some monocytes/macrophages, therapeutic manipulation of their activity may have undesirable consequences in pathological settings characterized by excessive angiogenesis, including cancer.^68,69^ Understanding how PAN mobilization and activity can be selectively regulated will therefore be essential.

Several limitations of this study warrant consideration. The murine hindlimb ischemia model represents an acute ischemic event and does not fully reproduce the chronic vascular pathology and inflammatory environment associated with peripheral arterial disease. Furthermore, although our human findings demonstrate a rapid ischemia-associated increase in circulating PANs, their anatomical origin and functional relevance remain unknown. Finally, our experiments do not establish whether the spleen exclusively stores PANs or additionally contributes to their phenotypic maintenance or differentiation. The functional equivalence of PANs originating from distinct anatomical reservoirs also remains to be determined.

In conclusion, our findings reveal a previously unrecognized mechanism by which the spleen contributes to vascular recovery following ischemic injury. Through sympathetic regulation of CXCL12, the spleen serves as a mobilizable reservoir of pro-angiogenic neutrophils that support restoration of tissue perfusion. The identification of a corresponding ischemia-responsive PAN population in humans suggests that this response may be conserved across species and highlights the potential of targeting neutrophil mobilization to enhance vascular repair.

## CONTRIBUTIONS

CAL, KP, CS, GC and RL designed, performed, and analyzed experiments. CAL, KP and MP wrote the manuscript. AS, HE and OP performed experiments. CFP and KM performed and designed human experiments. AW conceived and designed human experiments. MP conceived the study and designed and supervised experiments. All authors critically revised the manuscript.

## ACKNOWLEDGMENTS

The skillful support of BioVis Facility at Uppsala University is thankfully acknowledged. Emelie Öhnstedt is gratefully acknowledged for her role in developing the protocols for staining PANS in the spleen. This study was supported by the Swedish Research Council, Knut and Alice Wallenberg foundation, The Swedish Cancer Society, the Heart and Lung Foundation, and O. E. and Edla Johansson’s foundation.

---

**Supplemental figure 1.**
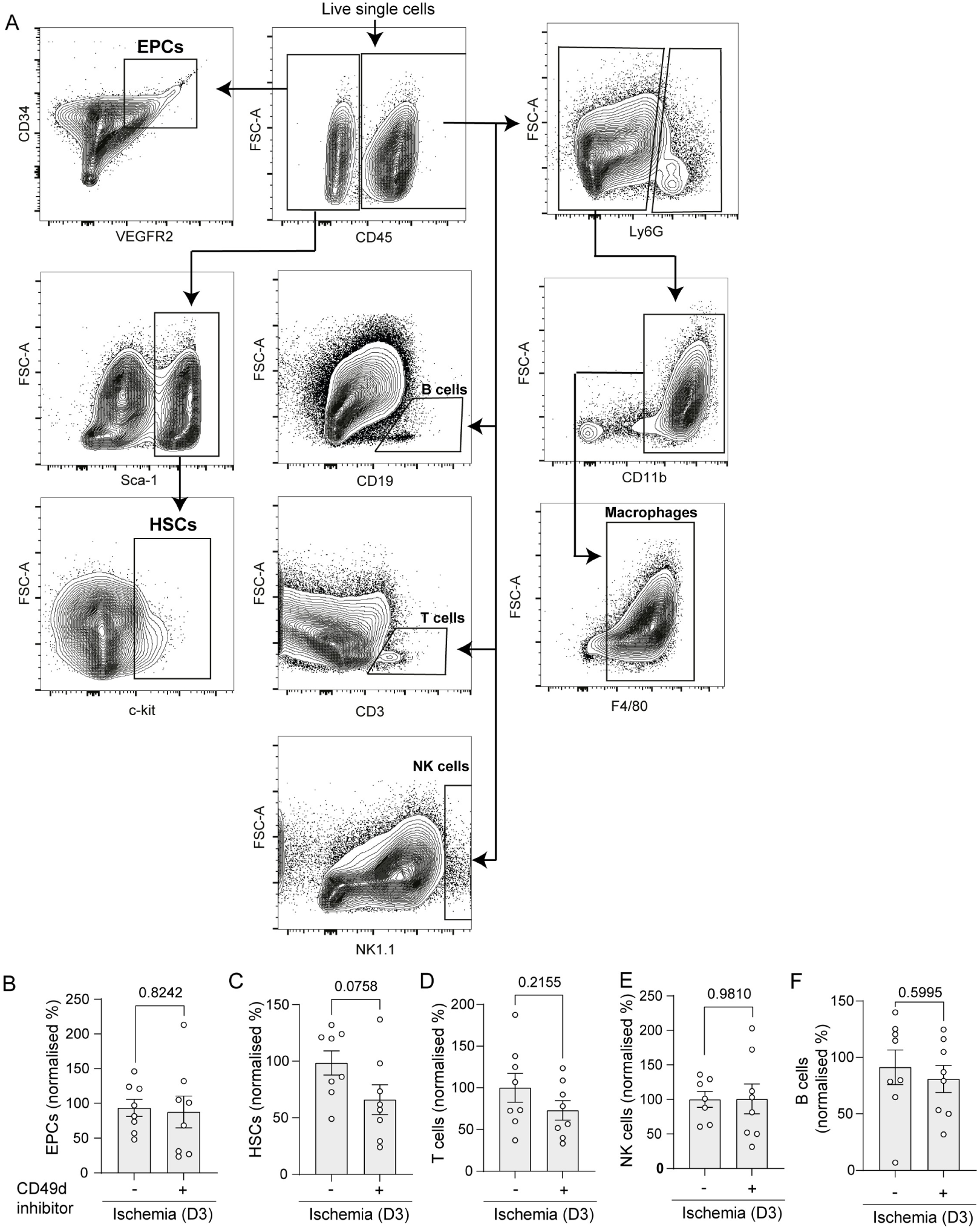
Representative gating strategies (A), and (B) normalised percentages of EPCs (B) HSCs (C), T cells (D), NK cells (E) and B cells (F) in CD49d inhibitor treated and control mice at 3 days post-HLI. Data are shown as average ± SEM. Unpaired students t-test (B-F).

**Supplemental Figure 2.**
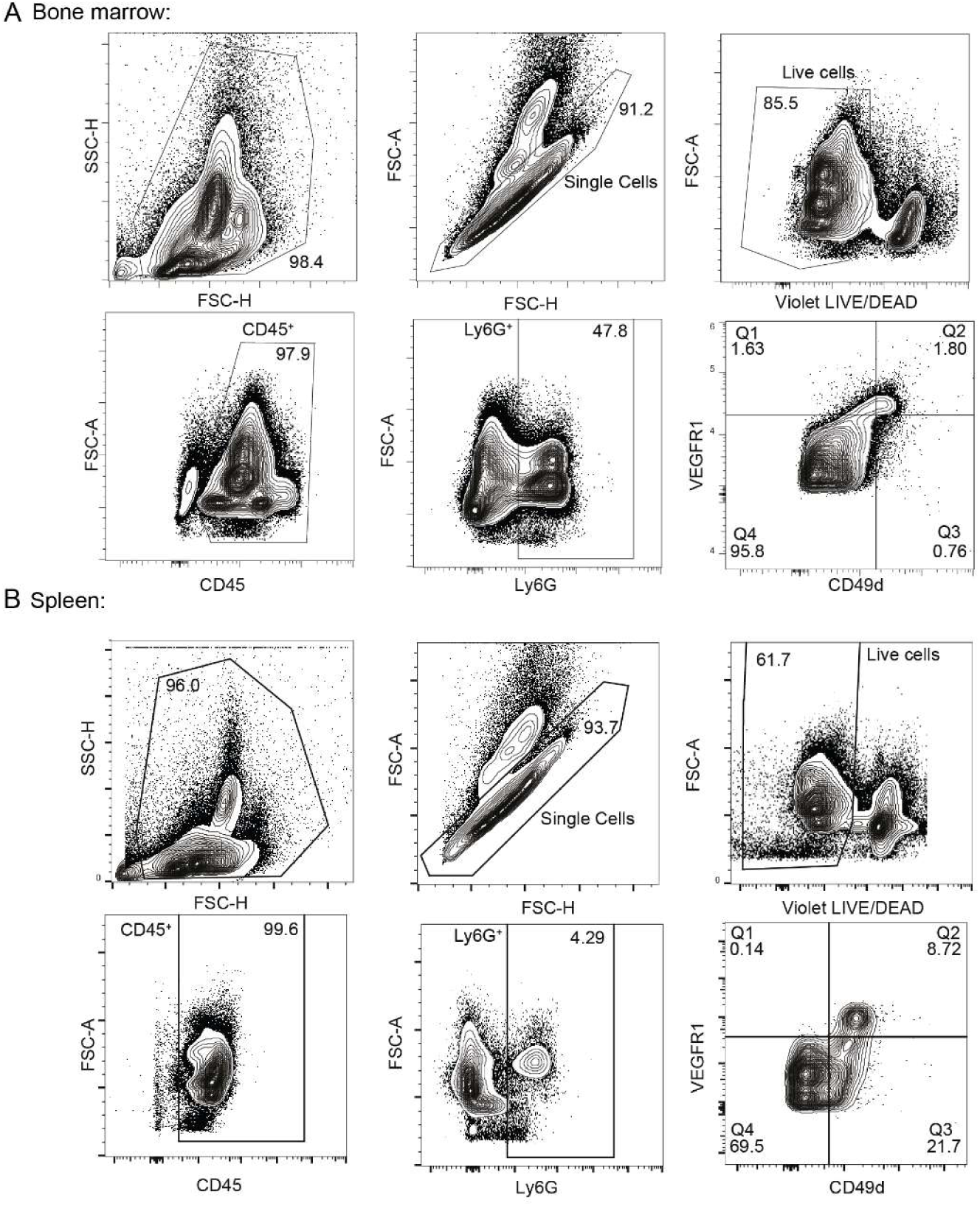
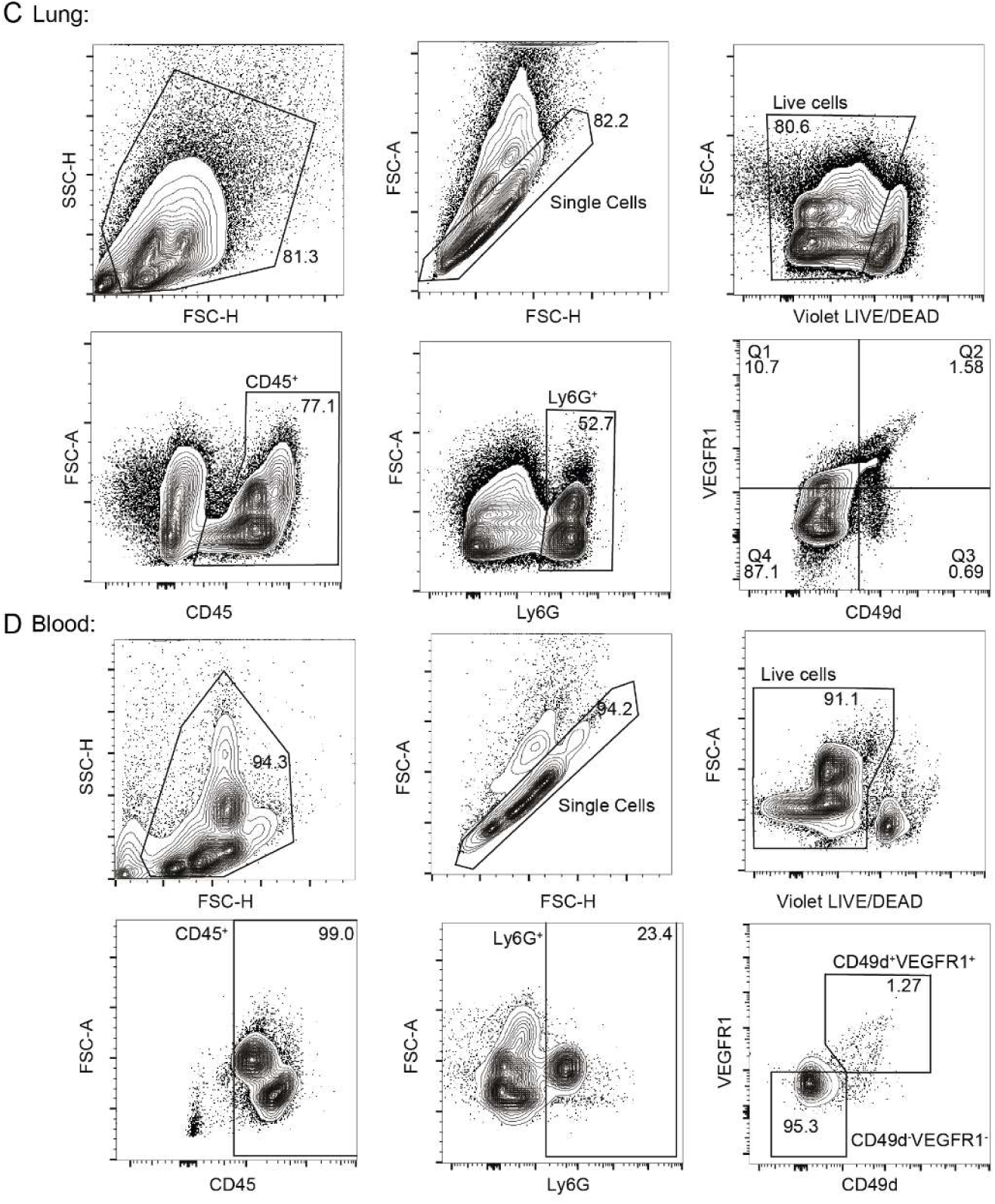
Representative gating strategies for classic neutrophils and PANs in the A) bone marrow. B) spleen, C) lung and D) blood.

**Supplemental Figure 3.**
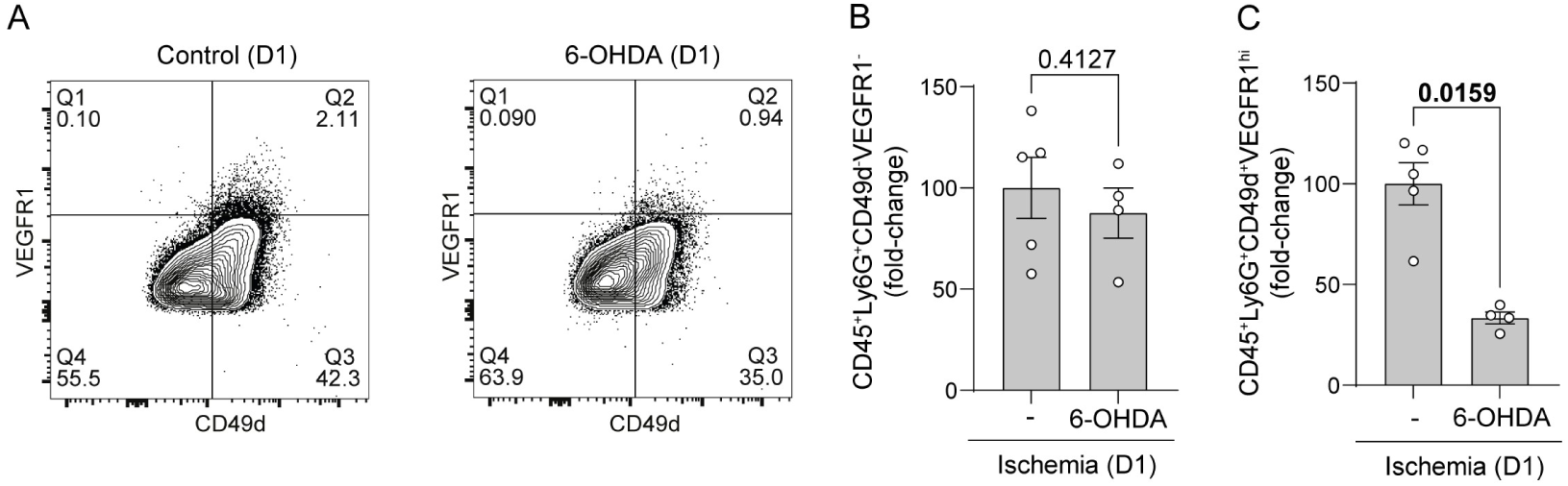
A) Representative contour plot of gating for classic neutrophils and PANs in the bone marrow of control (left) and 6-OHDA-treated (right) mice at 1 day post-HLI. Fold-change of C) classic neutrophils and D) PANs in the bone marrow of control and 6-OHDA-treated mice 1 day post-HLI. Mann-Whitney U test (B-C). Data are shown as average ± SEM.

**Supplemental Figure 4.**
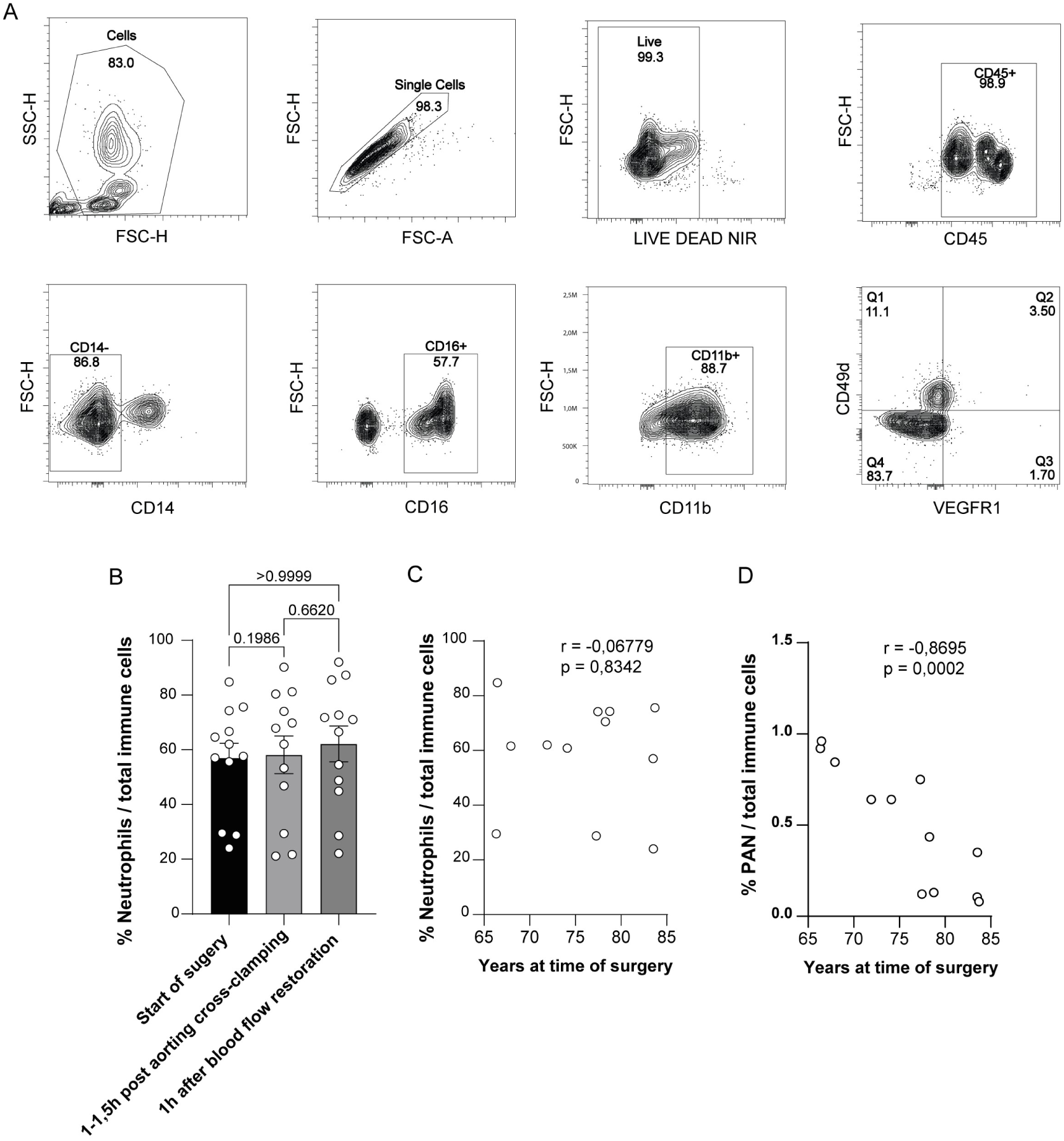
A) Representative contour plot of gating for human PANs in blood. B) Percentage of Neutrophils from total immune cells through surgical timepoints. C) Correlation between age and percentages of neutrophils from total immune cells. D) Correlation between age and percentage of PAN from total immune cells. Friedman test with Dunn’s multiple comparisons test (B). Data are shown as average ± SEM. Nonparametric Spearman correlation test (C-D).

## References

1. Kalogeris T, Baines CP, Krenz M, Korthuis RJ. Cell biology of ischemia/reperfusion injury. Int Rev Cell Mol Biol. 2012;298:229–317.

2. Kelly AG, Panigrahy D. Targeting Angiogenesis via Resolution of Inflammation. Cold Spring Harb Perspect Med. 2023;13:a041172.

3. Rizo-Téllez SA, Filep JG. Beyond host defense and tissue injury: the emerging role of neutrophils in tissue repair. Am J Physiol Cell Physiol. 2024;326:C661–C683.

4. Wang Y, Shou X, Wu Y, Li D. Immuno-inflammatory pathogenesis in ischemic heart disease: perception and knowledge for neutrophil recruitment. Front. Immunol. [Internet]. 2024 [cited 2026 Sept 3];15. Available from: https://www.frontiersin.org/journals/immunology/articles/10.3389/fimmu.2024.1411301/full

5. Heusch G. Myocardial ischemia/reperfusion: Translational pathophysiology of ischemic heart disease. Med [Internet]. 2024 [cited 2026 Sept 23];5:10–31. Available from: https://www.sciencedirect.com/science/article/pii/S2666634023004051

6. Roth GA, Mensah GA, Johnson CO, Addolorato G, Ammirati E, Baddour LM, Barengo NC, Beaton AZ, Benjamin EJ, Benziger CP, et al. Global Burden of Cardiovascular Diseases and Risk Factors, 1990–2019. JACC [Internet]. 2020 [cited 2026 Sept 23];76:2982–3021. Available from: 10.1016/j.jacc.2020.11.010

7. Saadeldeen AM, Mansour A, El-Dessouki AM, Fahim SA, Shaheen AM, Ismail RA, Salama RM, El-Shiekh RA, Khalifa HO. A comprehensive review of ischemic heart disease: pathophysiology, current treatments, natural products-based therapies, and nanotherapeutics. Front. Pharmacol. [Internet]. 2026 [cited 2026 Sept 23];17. Available from: https://www.frontiersin.org/journals/pharmacology/articles/10.3389/fphar.2026.1834343/full

8. Sabra M, Karbasiafshar C, Aboulgheit A, Raj S, Abid MR, Sellke FW. Clinical Application of Novel Therapies for Coronary Angiogenesis: Overview, Challenges, and Prospects. Int J Mol Sci [Internet]. 2021 [cited 2026 Sept 23];22:3722. Available from: https://pmc.ncbi.nlm.nih.gov/articles/PMC8038234/

9. Chen X, Yu W, Zhang J, Fan X, Liu X, Liu Q, Pan S, Dixon RAF, Li P, Yu P, et al. Therapeutic angiogenesis and tissue revascularization in ischemic vascular disease. J Biol Eng. 2023;17:13.

10. Kolaczkowska E, Kubes P. Neutrophil recruitment and function in health and inflammation. Nat Rev Immunol. 2013;13:159–175.

11. Rosales C. Neutrophils at the crossroads of innate and adaptive immunity. J Leukoc Biol. 2020;108:377–396.

12. Hidalgo A, Chilvers ER, Summers C, Koenderman L. The Neutrophil Life Cycle. Trends in Immunology. 2019;40:584–597.

13. Ley K, Laudanna C, Cybulsky MI, Nourshargh S. Getting to the site of inflammation: The leukocyte adhesion cascade updated. Nature Reviews Immunology. 2007;7:678–689.

14. Nourshargh S, Alon R. Leukocyte migration into inflamed tissues. Immunity. 2014;41:694–707.

15. Phillipson M, Kubes P. The Healing Power of Neutrophils. Trends in Immunology. 2019;40:635–647.

16. Mantovani A, Cassatella MA, Costantini C, Jaillon S. Neutrophils in the activation and regulation of innate and adaptive immunity. Nat Rev Immunol. 2011;11:519–531.

17. Sugimoto MA, Sousa LP, Pinho V, Perretti M, Teixeira MM. Resolution of Inflammation: What Controls Its Onset? Front Immunol. 2016;7:160.

18. Christoffersson G, Vägesjö E, Vandooren J, Liden M, Massena S, Reinert RB, Brissova M, Powers AC, Opdenakker G, Phillipson M. VEGF-Arecruits a proangiogenic MMP-9-delivering neutrophil subset that induces angiogenesis in transplanted hypoxic tissue. Blood. 2012;120:4653–4662.

19. Jaillon S, Ponzetta A, Di Mitri D, Santoni A, Bonecchi R, Mantovani A. Neutrophil diversity and plasticity in tumour progression and therapy. Nat Rev Cancer. 2020;20:485–503.

20. Massena S, Christoffersson G, Vägesjö E, Hidalgo CH, Giraud A, Lomei J, Westr S, Shibuya M, Claesson-welsh L, Welsh M, et al. Identification and characterization of VEGF-A – responsive neutrophils expressing CD49d, VEGFR1, and CXCR4 in mice and humans. Blood. 2015;126:2016–2027.

21. Piccard H, Muschel RJ, Opdenakker G. On the dual roles and polarized phenotypes of neutrophils in tumor development and progression. Critical Reviews in Oncology/Hematology. 2012;82:296–309.

22. Tazzyman S, Lewis CE, Murdoch C. Neutrophils: Key mediators of tumour angiogenesis. International Journal of Experimental Pathology. 2009;90:222–231.

23. Deniset JF, Kubes P. Recent advances in understanding neutrophils. F1000Res [Internet]. 2016 [cited 2026 Sept 3];5:2912. Available from: https://pmc.ncbi.nlm.nih.gov/articles/PMC5225409/

24. Jhunjhunwala S, Alvarez D, Aresta-DaSilva S, Tang K, Tang BC, Greiner DL, Newburger PE, von Andrian UH, Langer R, Anderson DG. Splenic progenitors aid in maintaining high neutrophil numbers at sites of sterile chronic inflammation. Journal of Leukocyte Biology. 2016;100:253–260.

25. Leuschner F, Rauch PJ, Ueno T, Gorbatov R, Marinelli B, Lee WW, Dutta P, Wei Y, Robbins C, Iwamoto Y, et al. Rapid monocyte kinetics in acute myocardial infarction are sustained by extramedullary monocytopoiesis. J Exp Med. 2012;209:123–137.

26. Swirski FK, Nahrendorf M. Cardioimmunology: the immune system in cardiac homeostasis and disease. Nat Rev Immunol. 2018;18:733–744.

27. Limbourg A, Korff T, Napp LC, Schaper W, Drexler H, Limbourg FP. Evaluation of postnatal arteriogenesis and angiogenesis in a mouse model of hind-limb ischemia. Nature Protocols. 2009;4:1737–1746.

28. Vågesjö E, Parv K, Ahl D, Seignez C, Herrera Hidalgo C, Giraud A, Amoêdo-Leite C, Korsgren O, Wallén H, Juusola G, et al. Perivascular Macrophages Regulate Blood Flow Following Tissue Damage. Circ Res. 2021;

29. Vasamsetti SB, Florentin J, Coppin E, Stiekema LCA, Zheng KH, Nisar MU, Sembrat J, Levinthal DJ, Rojas M, Stroes ESG, et al. Sympathetic Neuronal Activation Triggers Myeloid Progenitor Proliferation and Differentiation. Immunity. 2018;49:93–106.e7.

30. Zhang X, Lei B, Yuan Y, Zhang L, Hu L, Jin S, Kang B, Liao X, Sun W, Xu F, et al. Brain control of humoral immune responses amenable to behavioural modulation. Nature [Internet]. 2020 [cited 2025 Nov 19];581:204–208. Available from: https://www.nature.com/articles/s41586-020-2235-7

31. Bailey IB, Shaydakov ME, Shaw PM. Abdominal Aorta Aneurysm Repair [Internet]. In: StatPearls. Treasure Island (FL): StatPearls Publishing; 2026 [cited 2026 Sept 3]. Available from: http://www.ncbi.nlm.nih.gov/books/NBK430752/

32. Wanhainen A, Van Herzeele I, Bastos Goncalves F, Bellmunt Montoya S, Berard X, Boyle JR, D’Oria M, Prendes CF, Karkos CD, Kazimierczak A, et al. Editor’s Choice -- European Society for Vascular Surgery (ESVS) 2024 Clinical Practice Guidelines on the Management of Abdominal Aorto-Iliac Artery Aneurysms. Eur J Vasc Endovasc Surg. 2024;67:192–331.

33. Wanhainen A, Gombert A, Antoniou GA, Domingos LAF, Melo RG e, Grabenwöger M, Haulon S, Katsargyris A, Kölbel T, Loftus IM, et al. European Society for Vascular Surgery (ESVS) 2026 Clinical Practice Guidelines on the Management of Descending Thoracic and Thoraco-Abdominal Aortic Diseases – Editor’s Choice. European Journal of Vascular and Endovascular Surgery [Internet]. 2026 [cited 2026 Sept 3];71:172–270. Available from: https://www.ejves.com/article/S1078-5884(25)01319-X/fulltext

34. Chen J, Chemaly E, Liang L, Kho C, Lee A, Park J, Altman P, Schecter AD, Hajjar RJ, Tarzami ST. Effects of CXCR4 Gene Transfer on Cardiac Function After Ischemia-Reperfusion Injury. Am J Pathol [Internet]. 2010 [cited 2026 Sept 27];176:1705–1715. Available from: https://pmc.ncbi.nlm.nih.gov/articles/PMC2843462/

35. DeNies MS, Rosselli-Murai LK, Schnell S, Liu AP. Clathrin Heavy Chain Knockdown Impacts CXCR4 Signaling and Post-translational Modification. Front. Cell Dev. Biol. [Internet]. 2019 [cited 2026 Sept 27];7. Available from: https://www.frontiersin.org/journals/cell-and-developmental-biology/articles/10.3389/fcell.2019.00077/full

36. Kiani AA, Babaei F, Sedighi M, Soleimani A, Ahmadi K, Shahrokhi S, Anbari K, Nazari A. CXCR4 expression is associated with time–course permanent and temporary myocardial infarction in rats. Iran J Basic Med Sci [Internet]. 2017 [cited 2026 Sept 27];20:648–654. Available from: https://pmc.ncbi.nlm.nih.gov/articles/PMC5501928/

37. Christoffersson G, Lomei J, O’Callaghan P, Kreuger J, Engblom S, Phillipson M. Vascular sprouts induce local attraction of proangiogenic neutrophils. Journal of Leukocyte Biology. 2017;102:741–751.

38. Gerhardt T, Ley K. Monocyte trafficking across the vessel wall. Cardiovasc Res. 2015;107:321–330.

39. Medrano-Bosch M, Simón-Codina B, Jiménez W, Edelman ER, Melgar-Lesmes P. Monocyte-endothelial cell interactions in vascular and tissue remodeling. Front Immunol. 2023;14:1196033.

40. Aulakh GK. Neutrophils in the lung: “the first responders”. Cell Tissue Res. 2018;371:577–588.

41. Murray K, Godinez DR, Brust-Mascher I, Miller EN, Gareau MG, Reardon C. Neuroanatomy of the spleen: Mapping the relationship between sympathetic neurons and lymphocytes. PLOS ONE [Internet]. 2017 [cited 2026 Sept 28];12:e0182416. Available from: https://journals.plos.org/plosone/article?id=10.1371/journal.pone.0182416

42. Ding X, Wang H, Qian X, Cao Y, Wang Q, Ding X, Wang H, Qian X, Han X, Yang L, et al. Panicle-Shaped Sympathetic Architecture in the Spleen Parenchyma Modulates Antibacterial Innate Report Panicle-Shaped Sympathetic Architecture in the Spleen Parenchyma Modulates Antibacterial Innate Immunity. CellReports. 2019;27:3799–3807.e3.

43. Bellinger DL, Lorton D. Sympathetic Nerves and Innate Immune System in the Spleen: Implications of Impairment in HIV-1 and Relevant Models. Cells [Internet]. 2022 [cited 2026 Sept 28];11:673. Available from: https://www.mdpi.com/2073-4409/11/4/673

44. Capoccia BJ, Gregory AD, Link DC. Recruitment of the inflammatory subset of monocytes to sites of ischemia induces angiogenesis in a monocyte chemoattractant protein-1-dependent fashion. J Leukoc Biol [Internet]. 2008 [cited 2026 Oct 11];84:760– 768. Available from: https://pmc.ncbi.nlm.nih.gov/articles/PMC2516891/

45. Semerad CL, Liu F, Gregory AD, Stumpf K, Link DC. G-CSF is an essential regulator of neutrophil trafficking from the bone marrow to the blood. Immunity. 2002;17:413– 423.

46. Mebius RE, Kraal G. Structure and function of the spleen. Nat Rev Immunol. 2005;5:606–616.

47. Huang Y, Liu Z, Li M, Wang D, Ye J, Hu Q, Zhang Q, Lin Y, Chen R, Liang X, et al. Deciphering the impact of aging on splenic endothelial cell heterogeneity and immunosenescence through single-cell RNA sequencing analysis. Immun Ageing [Internet]. 2024 [cited 2026 Sept 28];21:48. Available from: 10.1186/s12979-024-00452-1

48. Richter M, Von Göwels J, Fähndrich M, Lipgens Fernandez C, Grohn K, Welzel M, Schwarz D, Lang A, Polzin A, Reinartz Groba S, et al. Depth of neutrophil mobilization stratifies survival in ST-elevation myocardial infarction. Nat Cardiovasc Res [Internet]. 2026 [cited 2026 Sept 28];5:693–704. Available from: https://www.nature.com/articles/s44161-026-00836-0

49. Terashima T, Wiggs B, English D, Hogg JC, Van Eeden SF. Polymorphonuclear leukocyte transit times in bone marrow during streptococcal pneumonia. American Journal of Physiology - Lung Cellular and Molecular Physiology. 1996;271.

50. Ballesteros I, Rubio-Ponce A, Genua M, Lusito E, Kwok I, Fernández-Calvo G, Khoyratty TE, van Grinsven E, González-Hernández S, Nicolás-Ávila JÁ, et al. Co-option of Neutrophil Fates by Tissue Environments. Cell. 2020;183:1282–1297.e18.

51. Caldwell CC, Kojima H, Lukashev D, Armstrong J, Farber M, Apasov SG, Sitkovsky M V. Differential Effects of Physiologically Relevant Hypoxic Conditions on T Lymphocyte Development and Effector Functions. The Journal of Immunology. 2001;167:6140–6149.

52. Schioppa T, Uranchimeg B, Saccani A, Biswas SK, Doni A, Rapisarda A, Bernasconi S, Saccani S, Nebuloni M, Vago L, et al. Regulation of the Chemokine Receptor CXCR4 by Hypoxia. Journal of Experimental Medicine. 2003;198:1391–1402.

53. Okuyama H, Krishnamachary B, Zhou YF, Nagasawa H, Bosch-Marce M, Semenza GL. Expression of vascular endothelial growth factor receptor 1 in bone marrow-derived mesenchymal cells is dependent on hypoxia-inducible factor 1. Journal of Biological Chemistry. 2006;281:15554–15563.

54. Ajmo CT, Collier LA, Leonardo CC, Hall AA, Green SM, Womble TA, Cuevas J, Willing AE, Pennypacker KR. Blockade of adrenoreceptors inhibits the splenic response to stroke. Experimental Neurology. 2009;218:47–55.

55. Scheiermann C, Kunisaki Y, Lucas D, Chow A, Jang JE, Zhang D, Hashimoto D, Merad M, Frenette PS. Adrenergic nerves govern circadian leukocyte recruitment to tissues. Immunity. 2012;37:290–301.

56. Ao T, Kikuta J, Sudo T, Uchida Y, Kobayashi K, Ishii M. Local sympathetic neurons promote neutrophil egress from the bone marrow at the onset of acute inflammation. Int Immunol. 2020;32:727–736.

57. Méndez-Ferrer S, Lucas D, Battista M, Frenette PS. Haematopoietic stem cell release is regulated by circadian oscillations. Nature. 2008;452:442–447.

58. De Filippo K, Rankin SM. CXCR4, the master regulator of neutrophil trafficking in homeostasis and disease. Eur J Clin Invest. 2018;48 Suppl 2:e12949.

59. Pillay J, Tregay N, Juzenaite G, Carlin LM, Pirillo C, Gaboriau DCA, Farahi N, Summers C, Lo Celso C, Chilvers ER, et al. Effect of the CXCR4 antagonist plerixafor on endogenous neutrophil dynamics in the bone marrow, lung and spleen. J Leukoc Biol. 2020;107:1175–1185.

60. Cyster JG. Chemokines and the Homing of Dendritic Cells to the T Cell Areas of Lymphoid Organs. J Exp Med [Internet]. 1999 [cited 2026 Oct 11];189:447–450. Available from: https://pmc.ncbi.nlm.nih.gov/articles/PMC2192905/

61. Katayama Y, Battista M, Kao W-M, Hidalgo A, Peired AJ, Thomas SA, Frenette PS. Signals from the sympathetic nervous system regulate hematopoietic stem cell egress from bone marrow. Cell. 2006;124:407–421.

62. Meloni A, Pistoia L, Ciancio A, Santamaria V, Sorrentino F, Cirotto C, Quota A, Romano N, Maggio A, Righi R, et al. Splenectomy Is a Risk Factor for Cardiac Complications in TM. Blood [Internet]. 2017 [cited 2026 May 28];130:2225. Available from: 10.1182/blood.V130.Suppl_1.2225.2225

63. Li Y, Stone JR. The impact of splenectomy on human coronary artery atherosclerosis and vascular macrophage distribution. Cardiovascular Pathology [Internet]. 2016 [cited 2026 May 28];25:453–460. Available from: https://www.sciencedirect.com/science/article/pii/S1054880716300710

64. Tsai M-S, Chou S-E, Lai H-S, Jeng L-B, Lin C-L, Kao C-H. Long-Term Risk of Acute Coronary Syndrome in Splenectomized Patients Due to Splenic Injury. Medicine (Baltimore) [Internet]. 2015 [cited 2026 May 28];94:e610. Available from: https://pmc.ncbi.nlm.nih.gov/articles/PMC4553951/

65. Vafadarnejad E, Rizzo G, Krampert L, Arampatzi P, Arias-Loza A-P, Nazzal Y, Rizakou A, Knochenhauer T, Bandi SR, Nugroho VA, et al. Dynamics of Cardiac Neutrophil Diversity in Murine Myocardial Infarction. Circ Res. 2020;127:e232–e249.

66. Amoedo-Leite C, Parv K, Testini C, Herrera-Hidalgo C, Xu F, Giraud A, Malaquias M, Fasterius E, Holl D, Seignez C, et al. Macrophages upregulate mural cell-like markers and support healing of ischemic injury by adopting functions important for vascular support. Nat Cardiovasc Res [Internet]. 2024 [cited 2025 Oct 1];3:685–700. Available from: https://www.nature.com/articles/s44161-024-00478-0

67. Patel AS, Ludwinski FE, Kerr A, Farkas S, Kapoor P, Bertolaccini L, Fernandes R, Jones PR, McLornan D, Livieratos L, et al. A subpopulation of tissue remodeling monocytes stimulates revascularization of the ischemic limb. Sci Transl Med. 2024;16:eadf0555.

68. Carmeliet P, Jain RK. Molecular mechanisms and clinical applications of angiogenesis. Nature. 2011;473:298–307.

69. Hanahan D, Weinberg RA. Hallmarks of cancer: the next generation. Cell. 2011;144:646–674.

